# In vivo imaging of autophagy dynamics in the intact mammalian brain using a fluorescence lifetime-based sensor

**DOI:** 10.64898/2025.12.25.693346

**Authors:** Maya Maman, Kota Ando, Natalie Fisher, Connon Thomas, Yoel Levy, Gregory Bond, Tomer Kagan, Nikol Malchenko, Muriel Desbois, Shelly Ben Zaken, Shay Orovan, Gal Uzan, Sharbel Eid, Guy Naftchi, Shireen Sarji, Michal Lapidot, Rebecca Rapaport, Emanuela Perah Break, Maayan Gal, Anza Darehshouri, Brock Grill, Yusuke Hirabayashi, Benjamin Scholl, Tal Laviv

## Abstract

Autophagy regulates neuronal development, plasticity, and metabolism, yet our ability to monitor its spatial and temporal organization within intact mammalian neural circuits remains limited. We developed an imaging approach combining a pH-sensitive fluorescence lifetime readout with independent morphological labeling to quantify autophagic vesicle state, abundance, and trafficking in vivo. Following pharmacological, genetic, and ultrastructural validations, we resolved compartment-specific autophagy levels across somata, dendrites, and axons of layer 2/3 cortical excitatory neurons in the intact mouse brain. We uncovered distinct patterns of dendritic autophagy in vivo, revealing faster autophagic vesicle trafficking in distal than proximal dendrites, that were preferentially sensitive to sensory experience. Using a newly generated transgenic mouse line, we found distinct cell-type-specific basal autophagy in excitatory neurons, interneurons, astrocytes, and oligodendrocytes. These findings reveal the spatial, cell-type-specific, and activity-dependent organization of autophagy in the brain and establish a framework for investigating its regulation during development, plasticity, and disease.

## Introduction

Homeostasis is critical for cellular and organismal survival, metabolism, and function, by maintenance of a stable intracellular milieu despite ongoing changes in the environment. One prominent pathway to maintain homeostasis is autophagy, an evolutionarily conserved catabolic process originally identified in yeast ^1^, in which proteins, lipids and organelles are degraded via the lysosomal pathway ^2,3^. Ever since its initial characterization, overarching evidence suggests autophagy plays an instrumental role in maintenance and development of the nervous system with important roles in synaptic function and plasticity ^4–6^. Numerous human neurodevelopmental diseases are associated with mutations in genes and proteins which orchestrate autophagy ^7–11^. Indeed, genetic manipulations of key autophagy related genes (ATGs), which initiate and regulate this complex pathway ^12^, impair proper formation of neuronal circuits ^4,13,14^. During aging, ongoing autophagy flux is necessary for maintaining neuronal health and autophagy dysfunction is associated with pathologies leading to neurodegeneration and toxic protein aggregations ^15–17^.

Despite this vital role, the spatial and temporal dynamics of autophagy in the intact mammalian brain remain poorly defined. Although numerous methods have been developed to monitor autophagy ^18^, their application has largely been restricted to cell-based assays, and ex vivo and in vitro preparations. Autophagy consists of the formation of the double-membrane autophagosome, followed by fusion with lysosomes to generate mature autolysosomes, whose acidic lumen supports cargo degradation. Widely used approaches to assess autophagic flux in living cells rely on fluorescent proteins that respond differently to the acidic environment of lysosomes. In particular, tandem fluorescent LC3 reporters exploit the differential pH sensitivity of GFP and RFP: GFP fluorescence is quenched in acidic compartments, whereas RFP remains detectable ^19,20^. Studies in both mammalian brain tissue ^21–23^ and neuronal cultures ^24,25^ have shown that autophagic vesicles in neuronal somata and dendrites are predominantly mature. However, existing fluorescence intensity-based methods primarily distinguish between immature autophagosomes (GFP/RFP-positive) and mature autolysosomes (RFP-only), providing a largely binary readout of compartment maturation. These detection criteria have limited the ability to distinguish subcellular compartmentalization and cell-type specific autophagy levels within the intact brain. Moreover, detecting and tracking autophagic vesicles within acidic organelles in dense brain tissue remains challenging for existing imaging approaches, largely restricting studies of autophagic vesicle trafficking in axons^24,25^ and dendrites ^26,27^ to neuronal cultures. The mammalian brain provides a unique environment for studying autophagy, distinguished from isolated cell-based systems by two key features. First, the brain maintains a tightly regulated metabolic environment that is relatively insulated from fluctuations in systemic nutrient availability. Second, because autophagic vesicle trafficking is sensitive to synaptic activity ^26–28^, the anatomical architecture of synaptic connectivity within neuronal circuits may shape autophagic dynamics. Accordingly, investigating autophagy in the intact brain requires efficient labeling of acidic organelles alongside visualization of neuronal morphology within dense and complex brain tissue.

Significant advances have recently been made towards development of new fluorescent proteins that show greater resilience in low pH ^29,30^. Katayama *et al* ^29^ designed TOLLES, a highly stable fluorescent protein in acidic environment. This protein was used as a potential Förster Resonance Energy Transfer (FRET) donor to YPet, a YFP variant. However, it was only used for ratiometric imaging, which allowed only a binary readout of mitophagy in fixed brain tissue ^29^. Fluorescence lifetime imaging microscopy (FLIM) has been the dominant approach to monitor protein signaling in brain slices ^31–33^ and in the intact brain^34–37^. The major advantages of FLIM for in vivo measurements include (a) more stringent and accurate quantification ^38^ (b) reduced dependence on absolute expression levels ^35,39^, and (c) the use of a single donor emission channel enabling simultaneous assessment of cellular morphology and function^40^.

Here, we describe a new imaging approach for monitoring autophagy in the intact mouse brain. We engineered a fluorescence lifetime-based pH sensor using a highly stable FRET donor with an independent morphological cell fill. We enable, for the first time, quantitative assessment of autophagy across cell types and their subcellular compartments in the mammalian brain and detect experience-dependent changes in autophagic dynamics.

## Results

### Engineering fluorescence lifetime-based method to quantitatively monitor autophagy state and dynamics in the mouse brain

Although optical reporters have enabled the visualization of autophagic vesicle maturation in cultured cells and fixed tissues, approaches for monitoring autophagy dynamics in the intact, living mammalian brain remain limited. In particular, quantitative measurements of autophagic maturation and vesicle trafficking at subcellular resolution pose substantial challenges in the dense tissue environment of the brain. To address this need, we set out to develop a fluorescence lifetime-based approach to monitor the pH changes accompanying autophagic vesicle maturation in vivo, while resolving their subcellular distribution and trafficking dynamics (**Fig. 1A**). A major advantage of FLIM is its robust standardization, which has enabled precise longitudinal monitoring of molecular signaling in the intact mammalian brain^34,35,37^. Importantly, FLIM requires detection of donor emission only, thereby allowing simultaneous labeling of neuronal morphology with red-shifted fluorophores ^40^, which is essential for resolving dynamics within dense brain tissue. We therefore first designed a FRET/FLIM-based approach to monitor pH changes associated with autophagic vesicle maturation. We began by examining the use of TOLLES, a highly pH tolerant fluorescent protein ^29^, as a possible FRET donor for FLIM. We performed two-photon FLIM (2pFLIM) of HEK cells expressing TOLLES and found that its fluorescence lifetime displays a mono-exponential decay of ∼3.15ns **(Fig. 1B, Extended Fig. 1A-B)**, making it a suitable FRET/FLIM donor. We then fused TOLLES with sREACh (sR), a dark YFP variant ^41^, previously used as a non-emitting FRET acceptor for FLIM based sensors ^31,36^. Expression of sR-TOLLES showed significant FRET, as indicated by a reduction in the donor fluorescence lifetime to 2.2ns **(Fig. 1B, Extended Fig. 1A-B)**. We hypothesized that FRET/FLIM could be used to quantitatively measure pH by leveraging pH-dependent changes in acceptor absorbance ^42^, which alter FRET efficiency and are consequently reflected in the fluorescence lifetime of TOLLES. To test this, we monitored fluorescence lifetime in HEK293 cells expressing TOLLES/sR-TOLLES, and used Nigericin-KCl to manipulate intracellular pH levels ^43^. We found that cells expressing TOLLES showed minimal changes in lifetime across pH levels, which is in line with the previous in vitro characterization ^29^ (**Fig. 1C, Extended Fig. 1C-D**). On the other hand, cells expressing sR-TOLLES showed pH-dependent changes in fluorescence lifetime, with a dynamic range of ∼0.8ns (**Fig. 1C, Extended Fig. 1C-D**). We further tested pH-dependent changes in purified proteins and found that the fluorescence lifetime of TOLLES was highly stable in neutral and acidic pH, and was not affected by the proteolytic enzyme Pepsin (**Extended Fig. 1E)**, in line with previous intensity characterization ^29^. On the other hand, sR-TOLLES showed short lifetime in neutral pH, which was elevated to long lifetime in both low pH with or without pepsin (**Extended Fig. 1E)**. sR-TOLLES lifetime was reversible in HEK cells upon incubation of the same cells in different pH levels, in comparison to stable lifetime of cells expressing TOLLES (**Extended Fig. 1F-I)**.

**Fig. 1.**
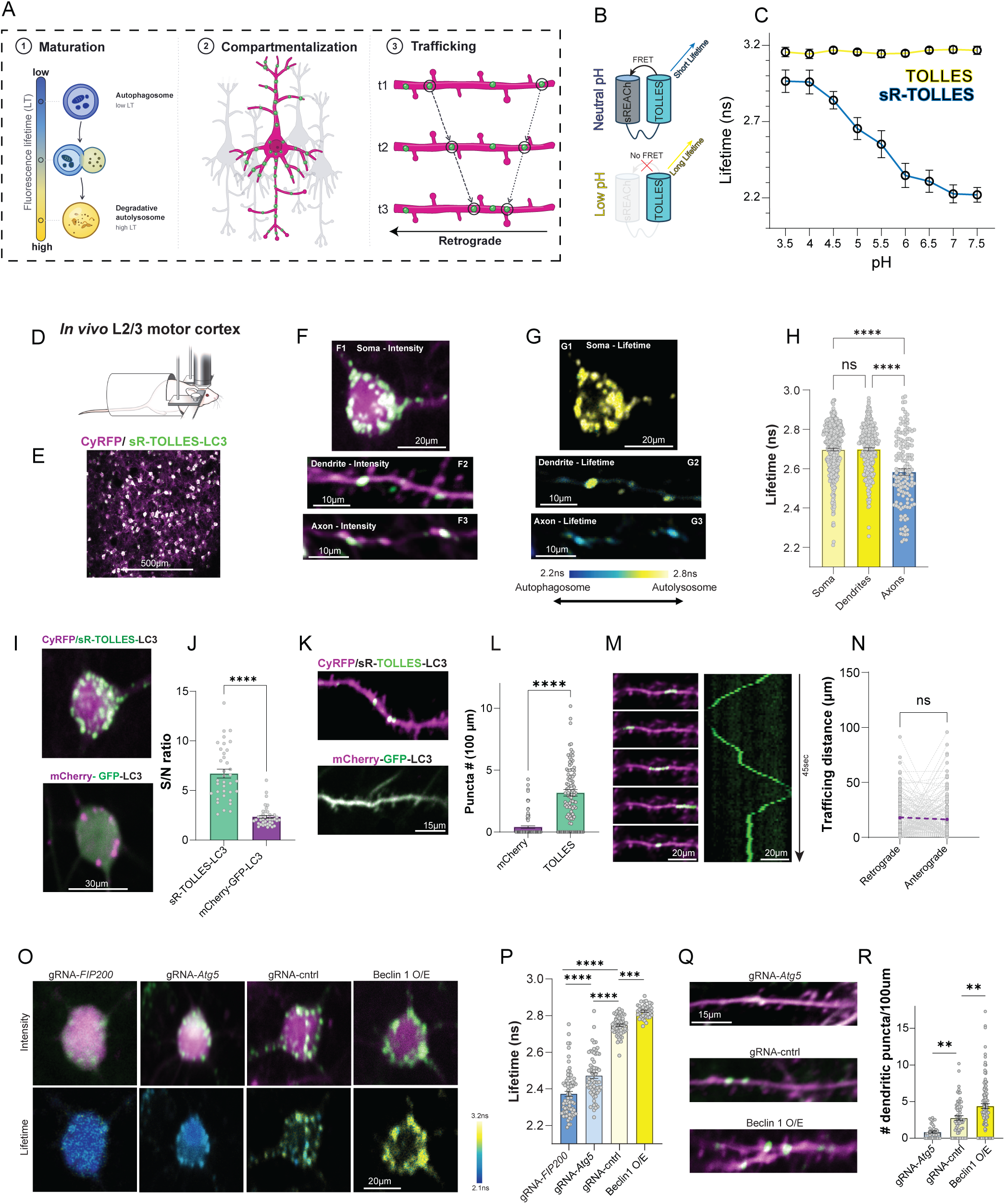
Engineering of a pH-dependent fluorescence lifetime biosensor for autophagy measurement in vivo. **(A)** Schematic illustration of the sensor readouts: 1-maturation of autophagic vesicles, 2-localization, and 3-trafficking, enabled by a red-shifted neuronal reporter. **(B)** Schematic illustration of the sREACh (sR)-TOLLES FRET pair in neutral and acidic pH. **(C)** pH dependent changes of mean fluorescence lifetime in cells expressing TOLLES (blue curve) and sR-TOLLES (yellow curve) (standard deviation bars). (**D**) Schematic illustration for *in vivo* imaging at adulthood **(E)** Representative field of view of L2/3 cortical neurons expressing sR-TOLLES-LC3 (green) and CyRFP (magenta). Scale bars-500µm **(F)** Fluorescence intensity representative images of L2/3 cell bodies (F1), dendrite (F2) and Axon (F3). Scale bars-20µm, 10µm, 10µm respectively. **(G)** Fluorescence lifetime representative images of L2/3 cell bodies (G1), dendrite (G2) and Axon (G3). Scale bars-20µm, 10µm, 10µm respectively. **(H)** Quantification of mean fluorescence lifetime in Soma (2.70±0.007ns, n=424 puncta), dendrites (2.70±0.008ns, n=271 puncta), and axons (2.58±0.016ns, n=143 puncta) in L2/3 neurons expressing the sensor. N=5 mice. p=0.9935 comparison between dendrites and soma. **(I)** Representative image of L2/3 cortical neurons expressing sR-TOLLES-LC3 and mCherry-GFP-LC3. Scale bar-30µm. **(J)** Quantification of signal to noise ratio (puncta/background) of sR-TOLLES-LC3 (6.71±0.45, n=36 cells) and mCherry-GFP-LC3 (2.34±0.16, n=40 cells), N=4 mice. **(K)** Representative fluorescence intensity image of a L2/3 dendrite expressing sR-TOLLES-LC3 (green) and CyRFP (magenta) (above) and mCherry-GFP-LC3 (magenta and green). Scale bar-15µm. **(L)** Quantification of LC3 positive puncta number per 100µm dendrite, in cells expressing mCherry-GFP-LC3 (0.4060 ± 0.0958 puncta/100µm, n = 95 dendritic branches) and sR-TOLLES-LC3 (3.171 ± 0.268 puncta/100µm, n = 98 dendritic branches) N=3-5 mice. **(M)** Representative maximum-intensity projection images (left) and corresponding kymograph (right) of a L2/3 dendrite expressing sR-TOLLES-LC3 (green) and CyRFP (magenta). The kymograph illustrates autophagic vesicle trafficking along the dendrite shown in the images, for a period of 45 seconds. Scale bars-20µm. **(N)** Quantification of paired retrograde and anterograde trafficking distance of autophagic vesicles along dendrites (retrograde: 17.86±1.313µm, Anterograde: 16.46±1.484µm, n=167 puncta) P=0.504. **(O)** Representative images showing sR-TOLLES/CyRFP fluorescence intensity and lifetime of L2/3 cell bodies expressing control gRNA, *FIP200* gRNA, *Atg5* gRNA, and over-expression of Beclin1. Scale bar −20µm. **(P)** Mean fluorescence lifetime in soma of *FIP200* gRNA (2.37±0.015ns, n=70 cells), *Atg5* gRNA (2.47±0.015ns, n=62 cells), control (2.75±0.007ns, n=59 cells) and Beclin1 Overexpression (2.82±0.006ns, n=64 cells) p=0.008. N=3-4 mice. **(Q)** Representative images showing sR-TOLLES/CyRFP fluorescence intensity and lifetime of L2/3 dendrites expressing control gRNA, *FIP200* gRNA, *Atg5* gRNA, and over-expression of Beclin1. Scale bar −15µm. **(R)** Quantification of the number of autophagic puncta detected in dendrites of *Atg5* KO (0.841±0.134 puncta/100µm, n=41 dendritic branches) p=0.0031 *Atg5* KO vs. control, control (2.767±0.306 puncta/100µm, n=61 dendritic branches), and Beclin O/E (4.386±0.033 puncta/100µm, n=112 dendritic branches). p=0.0015 beclin1 O/E vs. control. N=3-4 mice. Statistical difference was measured using one-way ANOVA followed by post-hoc Tukey’smultiple comparison test (H, P, R), unpaired two-tailed student t-test (J, L), and paired two-tailed student t-test (N). **** denotes p < 0.0001.

Next, we targeted the use of sR-TOLLES towards autophagy measurements. We generated a C-terminal fusion of *LC3B* with the sR-TOLLES FRET pair **(Extended Fig. 2A)**. We hypothesized that fluorescence lifetime changes would accompany acidification of autophagic compartments in a continuous, quantitative, and standardized manner. We first validated this approach in vitro, by transiently expressing sR-TOLLES-LC3 in mouse embryonic fibroblasts (MEF). Pharmacological induction of autophagy using the mTOR inhibitor Torin1 led to a robust increase in LC3 puncta that displayed longer fluorescence lifetime (**Extended Fig. 2B-C**), which was blocked by the addition of Bafilomycin, which mitigates lysosomal acidification (**Extended Fig. 2B-C)**. Importantly, MEFs devoid of *Atg5* ^44^, a core gene necessary for autophagy ^13,14^, did not change their lifetime upon induction of autophagy with Torin (**Extended Fig. 2B-C**), demonstrating the specificity of lifetime-based autophagy monitoring. We further tested if lifetime dependent changes correspond to gradual maturation of autophagic vesicles. We monitored fluorescence lifetime of sR-TOLLES-LC3 puncta and their colocalization with lysosomes, using a red-shifted LysoTracker. We found that the gradual increase in fluorescence lifetime of sR-TOLLES-LC3 following autophagy induction was associated with an increase in colocalization with lysosomes (**Extended Fig. 2D-G**).

To utilize in vivo 2pFLIM for monitoring autophagy in the intact mouse brain, we used in utero electroporation (IUE) to express sR-TOLLES-LC3 in layer 2/3 (L2/3) excitatory neurons. To morphologically label neurons, we combined CyRFP as a cell fill, due to its high cellular brightness and large Stokes shift ^40^, enabling efficient in vivo two-photon excitation with the same wavelength as TOLLES with distinct emission profiles. We performed cranial window surgery on adult mice, followed by in vivo 2pFLIM (**Fig. 1D**). This approach allows us to simultaneously visualize neuronal morphology and sR-TOLLES-LC3 puncta distribution, in individual cell bodies, dendrites and axons with subcellular resolution (**Fig. 1E-F**). Critically, we could quantify fluorescence lifetime for individual puncta in each neuronal compartment. We found a broad distribution of lifetime, in values that predominantly correspond to mature, acidic autophagic vesicles (**Fig. 1G-H**). Interestingly, we found significantly longer lifetime in somata and dendrites than in axons (**Fig. 1G-H**), suggesting that axons contain a higher proportion of autophagosomes, consistent with studies in neuronal cultures which found a higher autophagosome to autolysosome proportion in axons ^24,25^. Our findings reveal the spatial distribution of autophagic maturation in layer 2/3 cortical neurons of the intact adult mouse brain. We compared the fluorescence lifetime of sR-TOLLES-LC3 to that of TOLLES-LC3 which lacks a pH sensitive acceptor. We found that TOLLES-LC3 showed a longer lifetime in vivo, confirming the reliance of sR-TOLLES-LC3 on FRET (**Extended Fig. 3A-D**). Consistent with their mature state, sR-TOLLES-LC3 puncta also colocalized with lysosomal markers (Extended Fig. 3E-F).

Then, we compared the sensitivity of sR-TOLLES-LC3/CyRFP for in vivo imaging, with that of the canonical tandem mCherry-GFP-LC3. In vivo imaging of L2/3 cell bodies expressing mCherry-GFP-LC3 revealed diffuse cytosolic GFP fluorescence and distinct mCherry-positive puncta (**Fig. 1I, K**). This diffuse GFP distribution has previously been reported in neuronal cultures ^24,26^ and attributed to LC3 association with microtubules. Accordingly, the diffuse GFP signal makes it difficult to determine the mCherry/GFP ratio of individual puncta, limiting the use of this reporter to assess basal autophagy in the intact brain. Critically, neurons expressing sR-TOLLES-LC3/CyRFP exhibited markedly brighter puncta and a greater number of detectable structures (**Fig. 1I-L**), likely reflecting the higher tolerance of TOLLES to acidic environments compared with mCherry. This resulted in a significantly improved signal-to-noise ratio (S/N) relative to mCherry-GFP-LC3 (**Fig. 1I-J**). The advantage was even more pronounced in dendrites, where sR-TOLLES-LC3/CyRFP enabled clear visualization of individual puncta alongside dendritic morphology, whereas the low S/N of mCherry-GFP-LC3 limited the detection of individual puncta (**Fig. 1K-L**). This noticeable increase in signal to noise allowed us to readily visualize dynamic trafficking of sR-TOLLES-LC3 labeled puncta in vivo (**Fig. 1M, Extended Movie 1)**. The morphological cell-fill allowed us to develop a stringent analysis pipeline to calculate the speed and direction of individual puncta during in vivo imaging (**Extended Fig. 4**). We found that individual puncta show complex bi-directional movement in dendrites and within dendritic spines (**Fig. 1M**). On average, dendritic LC3 puncta displayed bi-directional anterograde or retrograde movement with no preferred orientation (**Fig. 1N**), which would indicate a potential role in regulation of local dendritic structure and function.

Lastly, we evaluated the sensitivity and specificity of sR-TOLLES-LC3 during in vivo imaging to endogenous autophagy levels. We used CRISPR/Cas9 to perturb endogenous autophagy by targeting *FIP200*, a critical protein for the formation of autophagosomes ^45^. Using IUE, we co-expressed Cas9/gRNA which targets *FIP200* (**Extended Fig. 5A-B**). L2/3 cells expressing Cas9/*FIP200*-gRNA alongside CyRFP/sR-TOLLES-LC3 showed a significantly shorter lifetime in the soma, with a marked reduction in somatic puncta compared to control cells (**Fig. 1O-P**). We further used CRISPR/Cas9 to reduce *Atg5* expression (**Extended Fig. 5C-E**), another critical upstream regulator of autophagy ^46^. L2/3 neurons expressing Cas9/gRNA targeting *Atg5* displayed a significantly shorter fluorescence lifetime of sR-TOLLES-LC3, compared to control conditions (**Fig. 1O-P**). Consistent with previous studies showing the accumulation of immature LC3 structures in cellular *Atg5* KO models^47^, our in vivo fluorescence lifetime analysis revealed that the remaining LC3 puncta in *Atg5*-deficient neurons less acidic (**Fig. 1O-P**). To test the ability to detect increased autophagy flux, we overexpressed Beclin1, previously reported to augment autophagy ^48^. L2/3 neurons overexpressing Beclin1 alongside sR-TOLLES-LC3/CyRFP showed significantly longer fluorescence lifetime compared to control conditions (**Fig. 1O-P**). Additionally, we found that dendrites of *Atg5* deficient neurons showed a robust decrease in the number of sR-TOLLES-LC3 puncta, whereas dendrites in cells overexpressing Beclin1 showed an increase in the number of autophagic puncta (**Fig. 1Q-R**).

Overall, our in vivo data demonstrate that this approach can quantitatively monitor basal autophagy levels in the intact mammalian brain with subcellular precision. We named this approach **FLApHY**, for **F**luorescence **L**ifetime based **A**utophagy **pH** anal**y**sis.

### Characterization and validation of FLApHY for in vivo imaging of autophagy

Fluorescence lifetime is a robust biophysical readout that is largely independent of fluorescence intensity. Accordingly, lifetime measurements of sR-TOLLES-LC3 puncta should enable reliable assessment of autophagic vesicle maturation. To test this, we correlated fluorescence lifetime measurements with ultrastructural characterization of the corresponding compartments using correlative light and electron microscopy (**CLEM; Fig. 2A–B)**. We first performed live imaging using 2pFLIM on MEFs expressing sR-TOLLES-LC3 following a 2h incubation with Torin to induce autophagy (**Fig. 2A1-A2**). Cells were then fixed and imaged by confocal microscopy to relocate the same cells for serial 3D SEM processing and imaging (**Fig. 2A3-A4**). This approach enabled us to identify autophagosomes (APs) and autolysosomes (ALs) within the same cell previously analyzed by live-cell 2pFLIM (**Fig. 2B**). Because of the relatively broad two-photon point-spread function, individual puncta could not be unambiguously assigned to single ultrastructural compartments. We therefore calculated the fluorescence lifetime within each region of interest (ROI) and compared it with the fraction of mature autophagic structures [AL/(AP+AL)] identified within the corresponding region in the SEM dataset (**Fig. 2C**). Fluorescence lifetime showed a significant positive correlation with the fraction of ALs **(Spearman r=0.65, p=0.02, n=16 ROIs**), demonstrating that fluorescence lifetime-based monitoring of sR-TOLLES-LC3 provides a quantitative approach for assessing autophagic vesicle maturation in living cells.

**Fig. 2.**
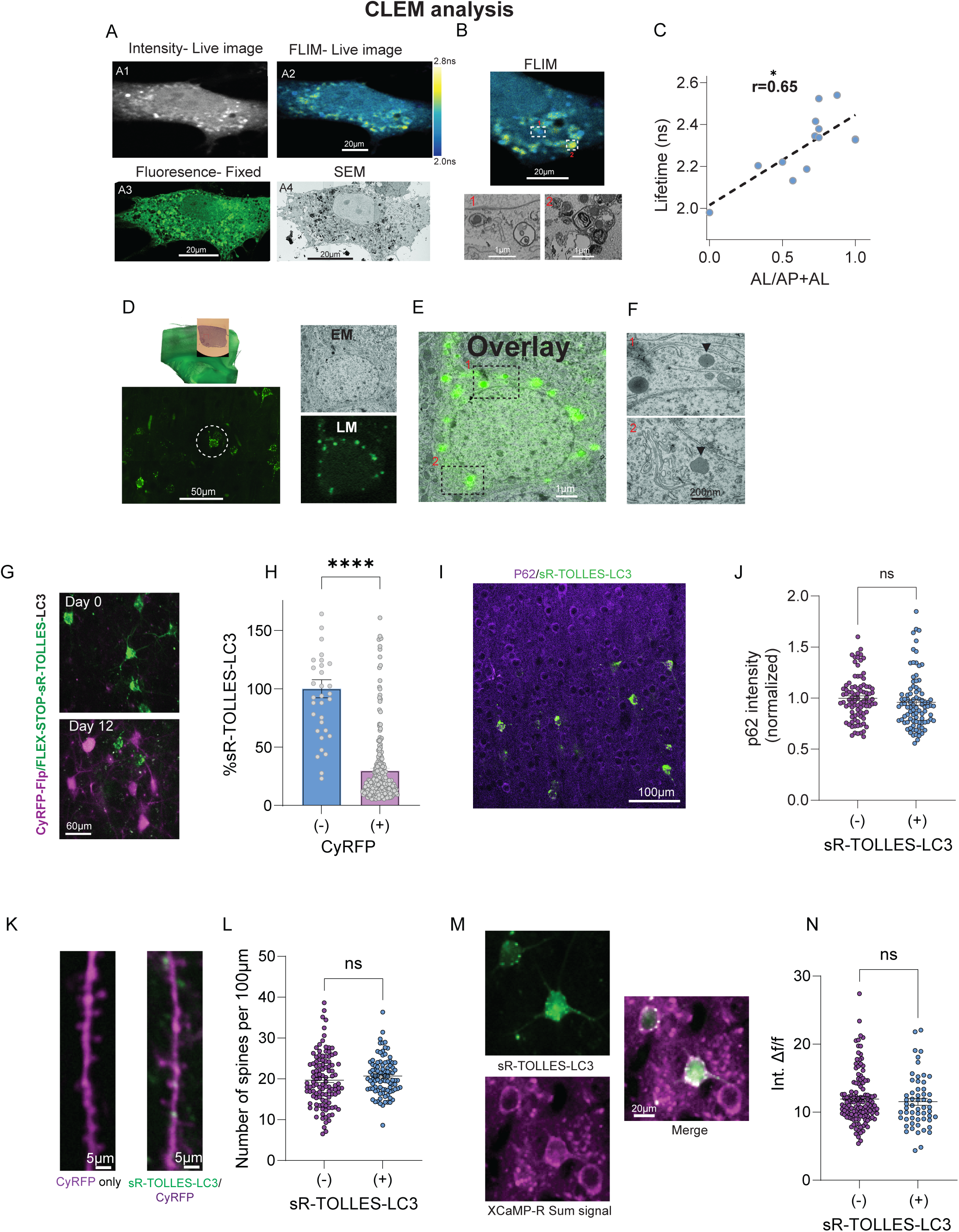
Structural and functional characterization of sR-TOLLES-LC3 expressing cells. **(A)** Representative whole-cell live images of TOLLES fluorescence intensity (**A1**) and fluorescence lifetime (**A2**) acquired by live-imaging following 2Hr of Torin application. Image of the same cell following fixation, followed by confocal microscopy (**A3**) and scanning electron microscopy (SEM; **A4**). Scale bars - 20µm **(B)** Representative correlation of individual puncta between FLIM (top) and SEM (bottom). ROI showing shorter lifetime (**1**) correlate with higher autophagosome/autolysosome in the SEM, whereas ROI (**2**) with longer lifetime, is correlated with higher autolysosmal distribution in SEM. Scale bars are 20µm for FLIM and 1µm for SEM. **(C)** Correlation between puncta fluorescence lifetime and lysosomal abundance within the correlated area. Spearman r=0.65, p=0.02. 16 ROIs. **(D)** Light microscopy (LM) field of view (left) and zoom-in (right) of a single cell expressing sR-TOLLES-LC3 in confocal light microscopy (LM) and electron microscopy (EM). **(E)** Overlay of EM and LM images of the same cell as in D **(F)** zoomed-in images, demonstrating TOLLES positive puncta colocalize with lysosomes. **(G)** Representative images of fields of view on the day of AAV-pSyp-Flp-P2A-Cre injection (Day 0 – sR-TOLLES-LC3 positive cells, green) and 12 days after injection (day 12 -not infected cells expressing sR-TOLLES-LC3, infected cells no longer express sR-TOLLES-LC3, now expressing CyRFP, magenta). Scale bar-60µm. **(H)** Quantification of the relative % of sR-TOLLES-LC3 signal, following 8-12 days in CyRFP positive (29.73±2.007%, n=235) and CyRFP negative cells (100±7.742%, n=34). N=4 mice. **(I)** Representative image of a fixed brain tissue slice expressing sR-TOLLES-LC3 (green) and stained for p62 (magenta). Scale bar-100µm. **(J)** Quantification of p62 levels, showing no difference between neurons expressing sR-TOLLES-LC3 and nearby non-expressing cells (sR-TOLLES-LC3: 0.9997 ± 0.023, n = 90 cells; nearby cells: 0.9591 ± 0.028, n = 90 cells). P= 0.2098 **(K)** Representative image and **(L)** quantification of spine number in dendritic branches expressing CyRFP alone (19.690±0.604, n=112 dendritic branches), or CyRFP together with sR-TOLLES-LC3 (20.680±0.466, n=95 dendritic branches) P=0.21. Scale bar-5µm. **(M)** Representative sum-intensity image, and **(N)** integrated ΔF/F quantification of XCaMP-R activity in cells expressing sR-TOLLES-LC3 (11.530±0.513, n=56) and nearby cells (11.950±0.331, n=129) P=0.4946. N=3 mice. Scale bar-20µm. Statistical difference was measured using unpaired two-tailed student t-test. **** denotes p < 0.0001.

To determine the subcellular localization of individual sR-TOLLES-LC3 puncta in the brain, we performed correlative light and electron microscopy (CLEM) in L2/3 neurons expressing the sensor (**Fig. 2D-F**). We first acquired high-resolution confocal images of chemically fixed cortical tissue and then used fiducial landmarks to relocate the same cells for EM imaging. Using this approach, we correlated fluorescence signals from 11 neuronal somata with their corresponding ultrastructural profiles and examined regions containing sR-TOLLES-LC3 puncta (**Fig. 2D-E**). The puncta corresponded to membrane-bound compartments containing electron-dense material and, in some cases, internal membranous structures (**Fig. 2F, Extended Data Table 1**). We further validated this by performing in vivo imaging of sR-TOLLES-LC3 alongside LAMP1-mScarlet, a lysosomal marker, and found predominant colocalization in the majority of somatic and dendritic puncta (**Extended Fig. 3E**). We also performed additional post-hoc staining of brain slices and found colocalization of sR-TOLLES-LC3 expression with Cathepsin D, a marker of functional lysosomes (**Extended Fig. 3F**). These findings are in line with our lifetime and structural analysis of the mature nature of somatic and dendritic puncta.

One potential concern in our experimental approach is that the stability of TOLLES, including its resistance to acidic pH and proteolysis, may hinder its clearance in vivo. To address this concern, we examined the turnover of sR-TOLLES-LC3 in vivo using a modified expression system in which production of new sR-TOLLES-LC3 is terminated by Cre recombinase, while expression of a reporter gene, CyRFP, is simultaneously activated by Flp recombinase (**Extended Fig. 6**). The corresponding plasmids were delivered by IUE, and adult mice were subsequently injected with an AAV expressing Flp-P2A-Cre to simultaneously activate CyRFP expression and terminate further sR-TOLLES-LC3 production. We then compared FLApHY expression before and following AAV injection. We found that sR-TOLLES-LC3 fluorescence was significantly reduced by 70% in CyRFP-positive neurons (**Fig. 2G-H**). Given the time required for Cre/Flp expression and recombination, these results indicate that sR-TOLLES-LC3 does not accumulate indefinitely in vivo and instead undergoes substantial turnover over a maximal period of days.

We further examined whether expression of sR-TOLLES-LC3 perturbs neuronal autophagy and physiology. Previous studies have shown that disruption of autophagy can produce measurable changes in neuronal structure ^49^ and activity ^50^. First, we used immunohistochemistry to measure p62, an established autophagic cargo that accumulates following inhibition of autophagic degradation. Neurons expressing sR-TOLLES-LC3 showed p62 levels comparable to those of neighboring non-expressing cells (**Fig. 2I-J**). The ability of FLApHY to be combined with a red-shifted reporter further allowed us to assess dendritic spine density (**Fig. 2K-L**) and spontaneous calcium activity using XCaMP-R ^51^ (**Fig. 2M-N**). In both analyses, sR-TOLLES-LC3-expressing L2/3 neurons showed no detectable differences compared with neighboring control neurons. Together, these results demonstrate that FLApHY enables quantitative analysis of basal autophagic states in vivo, undergoes substantial turnover rather than long-term accumulation, and produces no detectable effects on neuronal autophagy, morphology, or spontaneous activity.

### In vivo 2pFLIM autophagy measurements in *C. elegans*

We investigated whether FLApHY could be applied to other model organisms, focusing on *C. elegans*, in which autophagy plays an important role in development, survival, and longevity ^52,53^. We fused sR-TOLLES to LGG-1 (sR::TOLLES::LGG-1), the *C. elegans* homologue of LC3, and engineered transgenic *C. elegans* that overexpress this probe using a pan-neuronal promoter. This approach allowed us to quantify autophagy using fluorescence lifetime in the bundle of axons that forms the nerve ring (**Extended Fig. 7A**). Using this approach, we tested how impairing RPM-1, a negative regulator of autophagy initiation in axons ^54^, affects fluorescence lifetime of sR::TOLLES::LGG-1. A previous study found that *rpm-1* mutants display an increase in autophagosome formation in the axon bundle of the nerve ring ^54^. Using in vivo 2pFLIM, we found a significant decrease in the fluorescence lifetime of sR::TOLLES::LGG-1 in *rpm-1* mutants compared to wild-type animals (**Extended Fig. 7A-B**). This indicates that in *rpm-1* mutants there is an increase in lower pH autophagic vesicles in the nerve ring, which is likely to be due to an increase in autophagosomes. Thus, our findings are consistent with the prior study that observed increased formation of autophagosomes in the nerve ring of *rpm-1* mutants ^54^.

### Cell-specific regulation of basal autophagy levels in the mouse brain

Previous efforts to compare basal autophagy across cell types or brain regions have been hindered by the high basal autophagic flux in the adult brain. Existing imaging tools primarily classify compartments as mature or immature, limiting their ability to resolve differences among predominantly mature compartments. FLApHY’s unique advantages for in vivo imaging, namely high brightness and quantitative pH readout, prompted us to compare basal autophagy levels across distinct cell types in the intact mouse brain.

We generated a Cre-dependent knock-in (KI) mouse line by integrating a floxed FLApHY cassette into the Rosa26 locus under the control of a CAG promoter (**Extended Fig. 8A**). Homozygous FLApHY KI mice were healthy and fertile. We crossed these mice with CaMKII-Cre and CNP-Cre lines ^55^ to direct FLApHY expression to oligodendrocytes and excitatory neurons, respectively (Extended **Fig. 8B-C**). To label astrocytes, we combined PiggyBac transposition with IUE ^56^ resulting in sparse, bright expression of both CyRFP and sR-TOLLES-LC3 in individual cortical astrocytes (**Fig. 3A**). We compared lifetime measurements and subcellular distribution across these cell types and observed distinct basal autophagic profiles. CaMKII-positive cortical neurons predominantly contained acidic, mature puncta (**Fig. 3C-D**), consistent with our earlier findings in neurons sparsely labeled by IUE. In contrast, astrocytes exhibited shorter average lifetimes, with substantial variability among cells (**Fig. 3A, D**). Oligodendrocytes predominantly contained acidic, mature puncta, resembling neurons despite their shared glial lineage with astrocytes (**Fig. 3B, D**). Further in vivo analysis revealed compartment-specific differences within individual astrocytes: somatic puncta exhibited shorter lifetimes than those in astrocytic processes, suggesting a greater prevalence of immature autophagosomes in the soma (**Fig. 3E-G**). Autophagic vesicles were also localized to astrocytic end-feet (**Fig. 3F**). Unlike neuronal puncta, astrocytic puncta showed no detectable movement during imaging sessions. Oligodendrocytes, however, exhibited rapid, bidirectional movement of FLApHY puncta in vivo (**Extended Movie 2**).

**Fig. 3.**
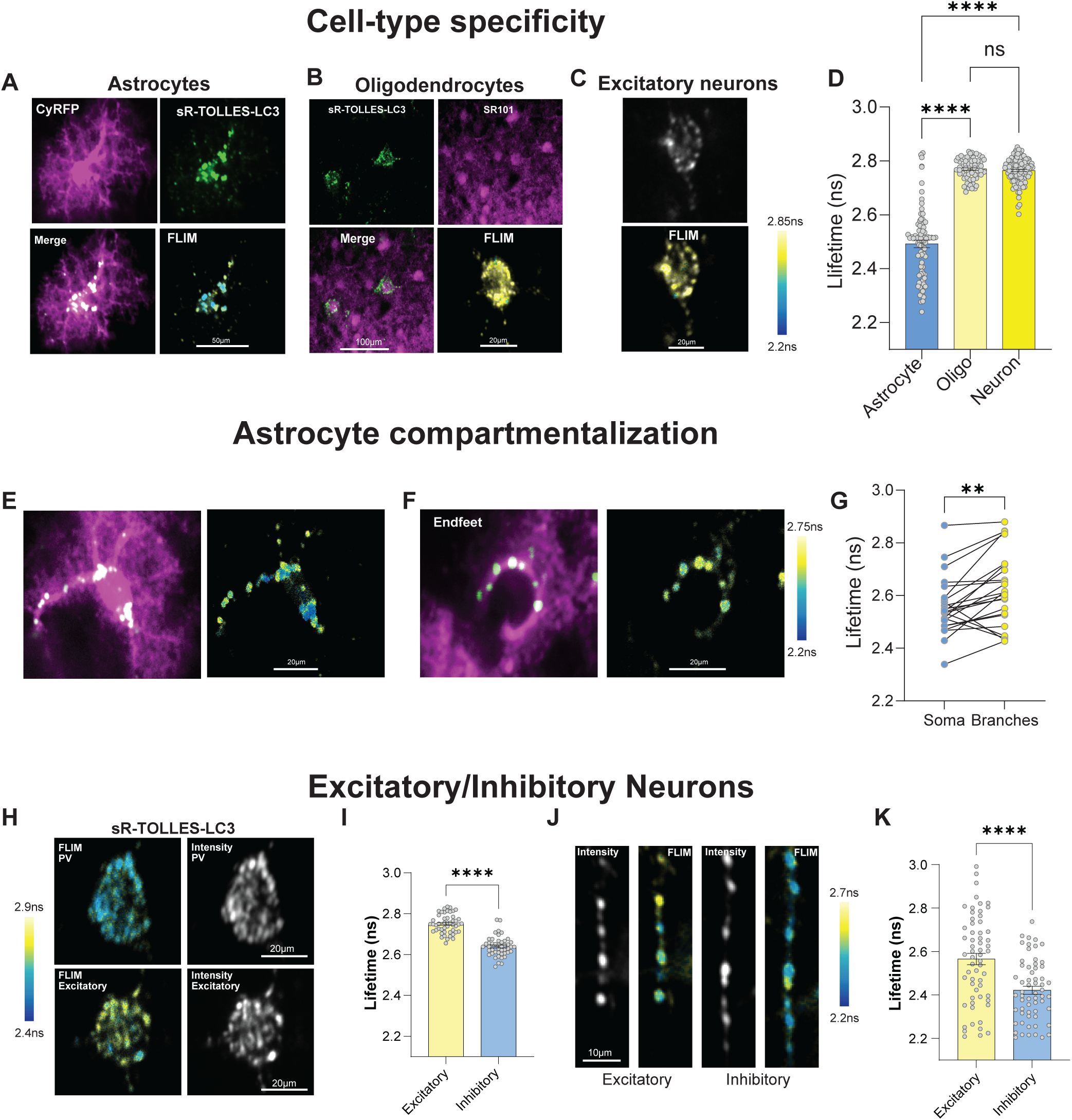
*In vivo* 2pFLIM unravels cell-type specific basal autophagy levels in the mouse brain. **(A)** Representative intensity and FLIM images of a cortical astrocytes expressing CyRFP (magenta) and sR-TOLLES-LC3 (green). Scale bar-50μm. **(B)** Representative intensity and FLIM images of oligodendrocytes expressing sR-TOLLES-LC3 in a transgenic FLEX-sR-TOLLES-LC3 /CNP-Cre mouse, labeled with SR101 (red). Scale bars −100µm and 20 µm respectively. **(C)** Representative intensity and FLIM images of cortical excitatory neuron in a transgenic sR-TOLLES-LC3-FLEX/CaMKIIa-Cre mouse. Scale bar - 20µm. **(D)** Quantification of TOLLES mean fluorescence lifetime in astrocytes (2.492±0.014ns, n=83), oligodendrocytes (2.770±0.005ns, n=61), and excitatory neurons (2.764±0.004ns, n=122). Oligodendrocytes versus Excitatory neurons: P=0.8903. N=3-5 mice. **(E)** Representative intensity (left) and FLIM (right) images of an astrocyte expressing sR-TOLLES-LC3 (green) and CyRFP (magenta). **(F)** Representative intensity (left) and FLIM (right) images of an astrocyte end-feet expressing sR-TOLLES-LC3 (green) and CyRFP (magenta). **(G)** Quantification of mean fluorescence lifetime in astrocytic cell body and branches. Each dot represents the average lifetime of all puncta within the cell body (2.558±0.023, n=23), paired with the corresponding average lifetime of puncta in the corresponding branches (2.623±0.029, n=23) from the same cell. P=0.0028. N=4 mice. **(H)** Representative fluorescence intensity and fluorescence lifetime images of excitatory L2/3 and PV positive cell bodies expressing sR-TOLLES-LC3. Scale bar-20µm. **(I)** Quantification of mean fluorescence lifetime of excitatory (2.75±0.007ns, n=48 cells) and inhibitory (2.64±0.007ns, n=46 cells) neurons. N=5 mice. **(J)** Representative fluorescence intensity and lifetime images of axons of excitatory and inhibitory neurons. Scale bar-10μm. **(K)** Quantification of Mean fluorescence lifetime of axonal vesicles in excitatory (2.56±0.026ns, n=60 puncta) and inhibitory (2.42±0.018ns, n=60 puncta) neurons. N=5 mice. Statistical difference was measured using one-way ANOVA followed by post-hoc Tukey’s multiple comparison test (D) and unpaired (I, K) and paired (G) two-tailed student t-test. **** denotes p < 0.0001.

Next, we set out to compare basal autophagy levels in different neuronal cell types. We injected an AAV with a Cre dependent sR-TOLLES-LC3 into a parvalbumin (PV)-Cre mouse line, and compared it to L2/3 excitatory neurons. Interestingly, we found that the fluorescence lifetime of PV interneuron cell bodies expressing sR-TOLLES-LC3 (**Extended Fig. 9)** was significantly shorter than that of excitatory cells (**Fig. 3H-I**). We observed the same difference for PV interneuron axons compared to L2/3 excitatory cell axons (**Fig. 3J-K**). These data suggest a higher proportion of autophagosomes in PV interneurons compared to L2/3 excitatory cells, which may be related to their unique function and metabolic needs^57,58^.

Overall, genetic labeling and the development of the FLApHY knock-in mouse enable systematic comparisons of autophagy across cell types in the intact brain. Our results reveal distinct patterns of autophagic maturation, subcellular compartmentalization, and vesicle trafficking, suggesting that autophagy is regulated in a cell-type-specific manner and may serve distinct roles in each cell type.

### Compartmentalized autophagy dynamics in dendrites

Previous studies have examined the regulation of autophagy in different neuronal subcellular compartments. Specifically, the majority of studies were conducted in murine neuronal cultures and Drosophila^28^, and focused on axonal autophagy, linking neuronal activity levels and the mobility of axonal autophagic vesicles ^24,28^. In addition, previous studies have found a role for dendritic autophagy during synaptic plasticity ^27,59^. However, the precise regulation and dynamics of dendritic autophagy within intact neuronal circuits remain unexplored. We used FLApHY to monitor autophagy in dendrites of L2/3 cells in the intact mouse motor cortex, by comparing the properties of autophagic vesicles in different dendritic compartments. We focused on basal-proximal dendrites and apical-distal dendrites (**Fig. 4A**), since a wealth of studies found distinct structural and functional organizational differences between these two compartments^60–62^. The comparison revealed significant differences: proximal dendrites showed marked increase in the density of autophagy vesicles compared to apical distal dendrites (**Fig. 4B-C**). The basal fluorescence lifetime of these populations was similar, indicating similar acidification and maturation state (**Fig. 4D**).

**Fig. 4.**
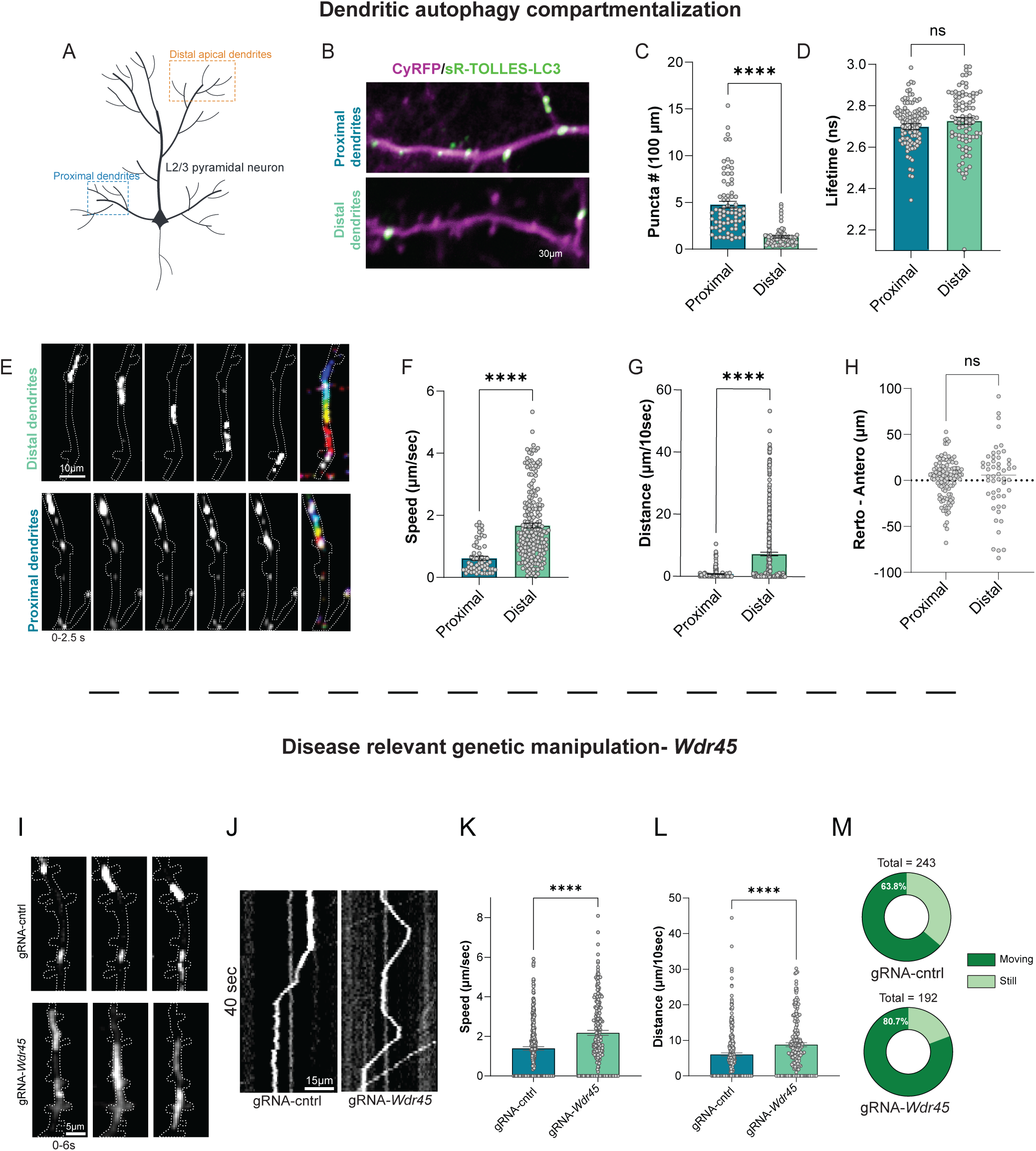
Compartmentalized dendritic autophagy dynamics in proximal and distal dendrites. **(A)** Schematic illustration of L2/3 pyramidal neuron, with proximal and distal dendrites. **(B)** Representative fluorescence intensity image of a L2/3 basal proximal and apical distal dendrites expressing sR-TOLLES-LC3 (green) and CyRFP (magenta) Scale bar-30µm. **(C)** Quantification of LC3 positive puncta number per 100µm dendrite, in proximal (4.773 ± 0.360, n = 76 dendritic branches) and distal (1.313 ± 0.114 puncta/100µm, n = 70 dendritic branches) dendrites. N=5 mice **(D)** Mean fluorescence lifetime of proximal (2.698 ± 0.0136ns, n = 105 puncta) and distal puncta (2.727 ± 0.0156ns, n = 89 puncta). p=0.1713. N=5 mice. **(E)** Representative maximal intensity image of autophagic vesicle movement in proximal and distal dendrites. The 16-color code bar represents 12.5 seconds total. **(F)** Quantification of the movement speed of individual autophagic vesicles in proximal (0.6200 ± 0.0651µm/sec, n = 56 puncta) and distal (1.671 ± 0.0791µm/sec, n = 204 puncta) branches. N=5 mice. **(G)** Quantification of the distance travelled over 10 seconds, by individual autophagic vesicles in proximal (1.659± 0.3292µm/10sec, n=137 puncta) and distal (7.299± 0.5801µm/10 sec, n=139 puncta) branches, including non-mobile puncta. N=5 mice. (**H**) Quantification of the overall distance travelled by individual puncta in proximal (1.363±1.980, n=115) and distal (1.477±5.113, n=52) dendritic branches, displays as total displacement in retrograde-anterograde direction. p=0.9799. (**I**) Representative maximal intensity images of sR-TOLLES-LC3 puncta movement in distal dendrites, in cells expressing control or *WDR45*-gRNA. (**J**) Representative kymographs of 40 seconds of distal dendrites with sR-TOLLES-LC3 positive puncta in cells expressing control or *WDR45*-gRNA. (**K**) Quantification of the movement speed of individual autophagic puncta in distal dendrites of Cas9/gRNA-control (1.39±0.091 µm/sec, n=243 puncta) and Cas9/gRNA-*WDR45* (2.18±0.125 µm/sec, n=192 puncta) cells. N=4 mice. (**L**) Quantification of the distance travelled over 10 seconds, by individual autophagic puncta in distal dendrites of Cas9/gRNA-control (6.00±0.461 µm/10sec. n=243 puncta) and Cas9/gRNA-*WDR45* (8.78±0.536 µm/10sec, n=192 puncta) cells. N=4 mice. (**M**) Comparisons of the ratio of moving dendritic puncta in Cas9/gRNA-control and Cas9/gRNA-*WDR45* cells. N=4 mice. Statistical difference was measured using unpaired two-tailed student t-test. **** denotes p < 0.0001.

Next, we tracked short-term mobility and dynamics of autophagic vesicles in anesthetized mice, in imaging sessions which lasted 50-60 seconds. We completed this analysis on 53 proximal basal dendrites and 177 apical distal dendrites. We found a dramatic difference in trafficking speed, where autophagic puncta in proximal dendrites were generally more stable and showed fewer mobile puncta than distal dendrites (**Fig. 4E-F**). Autophagic vesicles in distal compartments moved ∼3x faster than proximal ones (**Fig. 4E-F, Extended Movies 3 and 4**) and covered more distance (**Fig. 4G**). Both proximal and distal autophagic vesicles exhibit bidirectional trafficking, with no directional preference for anterograde or retrograde movement (**Fig. 4H**).

Finally, we sought to determine whether in vivo imaging of dendritic autophagy could reveal previously unrecognized phenotypes associated with genetic disorders of autophagy. We focused on *Wdr45*, which encodes WIPI4, a β-propeller domain-containing protein abundantly expressed in the brain. WIPI4 and related WIPI proteins play important roles in autophagy, and mutations in WDR45 are associated with severe forms of early-onset neurodegeneration ^63^. Although mouse models lacking Wdr45 exhibit profound structural and functional brain abnormalities, including axonal defects ^64^, whether these phenotypes arise directly from impaired neuronal autophagy remains unclear ^8,65^. To investigate the impact of *Wdr45* loss on neuronal autophagy in vivo, we used CRISPR/Cas9 to target *Wdr45* (**Extended Fig. 10A-C)** and co-expressed it with sR-TOLLES-LC3/CyRFP using IUE. We found that Wdr45-targeted neurons exhibited somatic sR-TOLLES-LC3 fluorescence lifetimes comparable to those of control neurons (**Extended Fig. 10D-E**). In contrast, imaging of distal dendrites revealed increased trafficking speed and travel distance of sR-TOLLES-LC3 puncta, along with a greater proportion of motile puncta (**Fig. 4I-M**), with no effect on overall puncta number (**Extended Fig. 10F**).

Overall, FLApHY enabled us to characterize autophagy levels and trafficking in dendrites in vivo, revealing distinct patterns of subcellular compartmentalization and their regulation by disease-associated genetic perturbations.

### Sensory deprivation leads to changes in dendritic autophagy

Lastly, we aimed to test the interplay between sensory experience and dendritic autophagy in the somatosensory cortex. Sensory experience profoundly modulates neuronal structure and function, in particular during early critical developmental periods ^66^. Prolonged whisker deprivation leads to long-lasting changes in synaptic structure and neuronal function^66–69^. However, the precise interplay between sensory experience and autophagy dynamics within the intact brain has not been explored.

To analyze individual dendrites with sufficient spatial resolution, we developed an AAV based approach to sparsely label individual cortical neurons, for simultaneous structural and autophagy measurements in vivo (**Fig. 5A**). We used a combination of AAVs encoding Cre-dependent sR-TOLLES-LC3, a Flp dependent AAV encoding CyRFP, and an AAV encoding Flp-P2A-Cre ^70^. Dilution of the Flp-Cre AAV led to sparsely labeled co-expressing neurons, ideal for dendritic in vivo imaging (**Fig. 5B**). We used AAV based sparse labeling of FLApHY and CyRFP to test the effect of sensory experience under two experimental settings: (1) Short-term sensory deprivation in adult mice (age >p45, 2 days trimming) (**Fig. 5C-D**) and (2) long-term sensory deprivation during the somatosensory critical period (age p28-p35, 14 days trimming) (**Fig. 5H-I**). We focused on dendritic trafficking in apical-distal dendrites, due to our earlier findings indicating they are highly dynamic. In short-term deprivation experiments, we compared control mice with mice which underwent contralateral whisker trimming. Our analysis reveals no significant change in the number of dendritic puncta, in the proportion of mobile puncta, or in trafficking speed (**Fig. 5E-G**). In contrast, we observed robust effects following long-term sensory deprivation. When compared to control littermates, mice that underwent long-term deprivation for 14 days showed significant reductions in the number of dendritic puncta (**Fig. 5J**). However, a greater proportion of these autophagic puncta were mobile (**Fig. 5K**) and displayed increased trafficking speed (**Fig. 5L**). In addition, we used FLApHY to analyze the lifetime of individual dendritic puncta, and found that long-term deprivation led to a significantly shorter lifetime (**Fig. 5M-N**). We conclude that long-term sensory deprivation altered autophagic levels, puncta distribution, and trafficking speed. Thus, the magnitude of sensory experience is likely a primary factor in dendritic autophagic vesicle formation and trafficking.

**Fig. 5.**
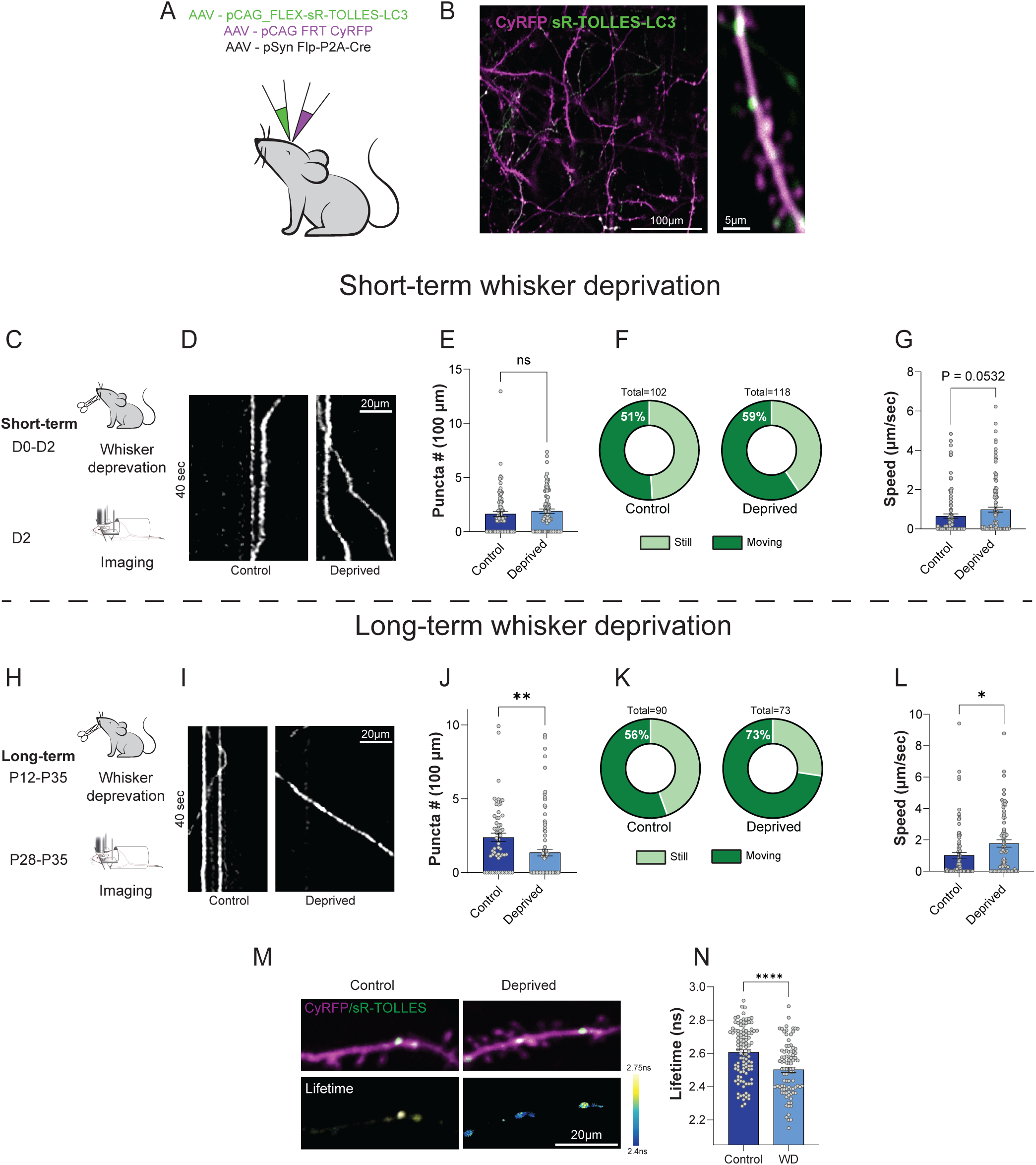
Sensory deprivation effects on dendritic autophagy in the somatosensory cortex. (**A**) Schematic illustration of AAV injections for sparse labeling of sR-TOLLES-LC3 and CyRFP (left) (**B**) representative images of distinct dendritic branches (right). Scale bars-100µm, 5µm respectively. (**C**) Experimental schematic illustration for *in vivo* imaging of individual distal dendrites expressing CyRFP and sR-TOLLES-LC3 in control and in mice that underwent short-term (48Hr) contralateral whisker deprivation. (**D**) Representative kymograph of individual puncta dendritic mobility during 40 sec. scale bar-20µm. (**E**) Comparison of LC3 positive puncta number per 100µm dendrite, in control (1.635 ± 0.2215 puncta/100µm, n = 83 dendritic branches) and short-term whisker deprived mice (1.891 ± 0.2085 puncta/100µm, n = 82 dendritic branches). p=0.40. N=5-6 mice. (**F**) Comparisons of the percentage of moving dendritic puncta in control and deprived mice. (**G**) Quantification of the dendritic movement speed of individual autophagic puncta in control (0.6513 ± 0.1096 µm/sec, n = 102 puncta) and short-term deprived mice (0.9822 ± 0.1268 µm/sec, n = 118 puncta). p=0.0532. N=5-6 mice. (**H**) Experimental schematic illustration for *in vivo* imaging of individual distal dendrites expressing CyRFP and sR-TOLLES-LC3 in control and in mice that underwent long-term contralateral whisker deprivation. (**I**) Representative kymograph of individual puncta dendritic mobility during 40 sec. scale bar-20µm for control and long-term deprived mice. (**J**) Comparison of LC3 positive puncta number per 100µm dendrite, in control (2.380 ± 0.2893 puncta/100µm, n = 59 dendritic branches) and long-term whisker deprived mice (1.351 ± 0.2325 puncta/100µm, n = 92 dendritic branches). p=0.0063. N=4 mice. (**K**) Comparisons of the percentage of moving dendritic puncta in control and long-term deprived mice. (**L**) Quantification of the dendritic movement speed of individual autophagic puncta in control (1.021 ± 0.1835 µm/sec, n = 86 puncta) and long-term deprived mice (1.771 ± 0.2319 µm/sec, n = 72 puncta). p=0.0111. N=4 mice. (**M**) Representative fluorescence intensity (top) images of sR-TOLLES-LC3 (green) and CyRFP (magenta) and fluorescence lifetime (bottom) of dendritic branches in control and whiskers deprived mice. Scale bar-20µm (**N**) Quantification of fluorescence lifetime in puncta located in dendritic branches of control (2.606±0.016, n=102) and long-term whiskers deprived (2.502±0.017, n=91) mice. N=4 mice. Statistical difference was measured using unpaired two-tailed student t-test. **** denotes p < 0.0001.

## Discussion

Here, we introduce FLApHY, a fluorescence-lifetime-based approach for quantifying autophagic vesicle maturation, compartmentalization, and trafficking in the intact mouse brain. FLApHY enables standardized comparisons across cell types and subcellular compartments, addressing a key challenge in studying autophagy in complex tissues.

FLApHY uses 2pFLIM to measure pH-dependent changes in FRET between the pH-insensitive donor (TOLLES) and the pH-sensitive dark acceptor (sREACh). Compared with intensity-based measurements, fluorescence lifetime is less sensitive to expression levels and variations in excitation and signal collection during in vivo imaging. Donor-only detection also permits simultaneous imaging with red-shifted markers of cellular structure or function, enabling localization and tracking of autophagic vesicles. Using FLApHY, we identified distinct autophagic profiles across neuronal cell types, astrocytes, and oligodendrocytes, as well as differences in autophagy levels and vesicle trafficking between proximal and distal dendrites. Distal dendrites exhibited greater vesicle motility, and sensory experience regulated vesicle trafficking in vivo. These findings reveal previously inaccessible features of autophagy organization and regulation in the mammalian brain.

### Potential limitations of this approach

One potential limitation of FLApHY is prolonged accumulation of the biosensor in lysosomes due to the pH resilience of TOLLES. We examined this possibility using several complementary approaches. Perturbation of the upstream autophagy regulators *Atg5* and *FIP200* reduced both sR-TOLLES-LC3 puncta abundance and fluorescence lifetime, consistent with reduced delivery of the sensor to acidic autophagic compartments. Conversely, augmentation of autophagy by Beclin1 overexpression increased fluorescence lifetime and dendritic puncta abundance. These results demonstrate that FLApHY remains sensitive to changes in endogenous autophagy despite biosensor overexpression. To assess reporter localization and persistence more directly, we combined CLEM analysis with an expression system that terminates new sR-TOLLES-LC3 production in vivo. CLEM showed predominant localization within autophagic compartments, while shutting off sensor transcription revealed significant turnover of existing reporter in vivo, indicating limited persistence. Together, these findings argue against persistent lysosomal accumulation dominating the readout. Nevertheless, sufficient biosensor expression is necessary to achieve reliable signal-to-noise ratios, and potential effects on endogenous autophagy should be assessed when applying FLApHY to new experimental systems.

### Dynamics and compartmentalization of dendritic autophagy

FLApHY’s sensitivity and compatibility with labeling of neuronal morphology enabled us to quantify and track dendritic autophagic puncta in the intact mouse brain for the first time. In L2/3 excitatory neurons, autophagic vesicles were more abundant in proximal dendrites, whereas those in distal dendrites moved faster and travelled greater distances. Previous studies in cultured neurons demonstrated sensitivity of dendritic autophagy to neuronal activity ^26^, but the random synaptic connectivity in neuronal cultures limits the analysis of dendritic compartments with defined anatomical functions. We examined how sensory experience regulates dendritic autophagy. At the Drosophila neuromuscular junction, starvation or synaptic stimulation mobilizes autophagic vesicles to axonal boutons ^28^. Here, we found that in the mouse somatosensory cortex, persistent sensory deprivation reduced dendritic autophagic puncta abundance while increasing trafficking speed and shortening fluorescence lifetime. Future studies could use FLApHY to enable further investigation of how these dynamics relate to structural and functional synaptic plasticity in vivo.

### Regulation of cell-type-specific autophagy in the brain

Quantitative lifetime measurements within mature acidic compartments enabled us to directly compare basal autophagy across cell types in the mouse cortex. Fluorescence lifetimes differed markedly between excitatory neurons and PV interneurons, suggesting reduced maturation of autophagic puncta, and potentially a higher fraction of autophagosomes, in PV interneuron somata and axons. Recent studies have established essential roles for autophagy in PV interneuron survival and brain plasticity ^71^. Future work could examine how their autophagic state relates to high firing rates and associated energetic and metabolic demands ^57^. We extended our analysis to cortical astrocytes, whose autophagy has been studied in culture ^72,73^ but remains poorly characterized in vivo. Astrocytes exhibited shorter fluorescence lifetimes than neurons, suggesting a higher fraction of immature autophagosomes. Within astrocytes, LC3 puncta in processes had longer lifetimes than those in the soma, indicating subcellular differences in vesicle maturation. These observations provide a basis for investigating the organization of astrocytic autophagy and its responses to genetic or environmental perturbations. To facilitate cell-type and brain-region specific studies, we generated a Cre-dependent FLApHY knock-in mouse line (**Extended Fig. 8A)** and used an oligodendrocyte Cre driver to examine autophagy in mature oligodendrocytes. Although previous studies have identified developmental roles for oligodendrocyte autophagy ^74,75^, its basal state and dynamics in vivo remain less understood. Oligodendrocyte fluorescence lifetimes resembled those of excitatory neurons and differed markedly from astrocytes, despite their shared glial classification. LC3 vesicles also exhibited rapid bidirectional motility (**Extended Movie 2**), highlighting the dynamic nature of autophagy in these cells. Together, these findings establish FLApHY as a platform for investigating cell-specific autophagy in brain function and homeostasis.

## Online Methods

### Mice

Animal experiments were approved by the Institutional Animal Care and Use Committee in Tel Aviv University.

The CAG-FLEX-STOP-FLApHY mice were produced by TransViragen (Chapel Hill, NC, USA). A Cas9 guide RNA (gRNA) targeting the mouse Gt(ROSA)26 1st intron (protospacer sequence 5’-GGAGTTGCAGATCACGA −3’) was cloned into a T7 promoter-sgRNA scaffold vector (TransViragen/UNC Chapel Hill Animal Models Core) and produced by T7 in vitro transcription (HiScribe T7 High Yield RNA Synthesis Kit, New England BioLabs), with RNeasy spin column purification (Qiagen) and elution in microinjection buffer (5 mM Tris-HCl pH 7.5, 0.1 mM EDTA). C57BL/6J zygotes were microinjected with 200 nM SpCas9 protein, 12.5 ng/µl gRNA and 10 ng/µl supercoiled donor vector CAG-FLEX-STOP-FLApHY. Microinjected embryos were implanted in recipient pseudopregnant B6D2F1/J females. Resulting pups were screened by PCR for the correct transgene integration. A founder harboring the transgene was mated to C57BL/6J females for germline transmission of the transgene.

In Utero Electroporation (IUE) was performed on timed-pregnant ICR mice (Envigo). For targeting of PV cells, we crossed ICR females with homozygous PV-Cre male mice (Stock No. 017320, The Jackson Laboratory) and heterozygous mice were used for experiments. CaMKIIα-Cre (Stock No. 005359, The Jackson Laboratory) and CNP-Cre ^55^ (Generous gift from Dr. Klaus Nave) were crossed with CAG-FLEX-STOP-FLApHY and heterozygous mice were used for experiments.

Both male and female mice were used throughout experiments. All animals were kept in a normal light/dark cycle (12 h/12 h, lights on at 7AM) and had free access to food and water. Mice were kept at 21±2°C, and the air change rate was 15-20 times per hour, with no humidity control.

### Plasmids and AAV construction

The sequence for TOLLES was kindly provided by Atsushi Miyawaki and the RIKEN DNA bank (RDB18222). sREACh-TOLLES fusion protein was prepared by adding a linker of ELGGGGSGGGGSGGGGSLE between the two fluorescent proteins. sREACh-TOLLES-LC3 was prepared by adding the *LC3B* sequence (From EGFP-LC3, a gift from Karla Kirkegaard, Addgene plasmid # 11546) preceded by a SGLRSA linker. The sREACh-TOLLES-LC3B coding sequence was then subcloned into CMV, CAG and AAV-CAG-FLEX backbones. PiggyBac vectors were used to express CyRFP, or sREACh-TOLLES-LC3B under the CAG promoter. AAV-pSyn-Flp-P2A-Cre was prepared by inserting the Flpe-P2A-Cre ^70^ sequence (kindly provided by Y. Voziyanov) into a AAV-pSynapsin backbone. AAV-FLEX-mCherry-GFP-LC3 was prepared by inserting mCherry-GFP-LC3 (Addgene #110060) into an AAV-CAG-FLEX backbone. The C*. elegans* autophagy reporter was prepared by replacing *LC3B* with the worm homologue *LGG1*. CAG-Lamp1-mScarlet was prepared from Addgene #98827. Beclin1 was a gift from Johan Jakobsson (Addgene plasmid # 107937). AAV-pCAG-FRT-CyRFP-FRT was a gift by David Fitzpatrick, CAG-CyRFP was from Addgene # 84356.

The autophagy sensor was used under the CMV promoter (mEGFP-C1 backbone) for in vitro experiments and expressed under the CAG promoter for IUE in mice. For expression in astrocytes, the piggyBac (PB) transposon system was used together with pCAG*-*hyPBase.

AAVs were packaged in AAV serotype 9 by the ELSC vector core facility. For gene editing via CRISPR/Cas9, we used the PX330 backbone (a gift from Feng Zhang, Addgene plasmid 42230). We introduced gRNA sequences according to target gene (5’-3’): *Atg5* (AAGATGTGCTTCGAGATGTG), *FIP200* (CTCAGTGTCAAATGTCAGCG), *WDR45* (GTCTTGGTTGAAATGTAGGC).

For the C. elegans experiment, sREACh-TOLLES was codon-optimized using the IDT website and fused with LGG1 gDNA (pBG-467). The sequence was then amplified and cloned into the pCR8 vector (pBG-GY1116). After nanopore sequencing (Plasmidsaurus), gateway LR recombination was used to create the final plasmid with the Prgef-1 promoter (Prgef-1::sR::Tolles::LGG-1 (pBG-GY1117). Plasmids were constructed using standard molecular biology methods including Polymerase chain reaction, Gibson assembly, enzyme restriction digestions and Q5 Site-Directed Mutagenesis (NEB). All plasmids were verified using Sanger Sequencing.

### Cell culture

HEK293T cells (ATCC) in passage number 12-20 were cultured in DMEM supplemented with 10% FBS, 1% L-Glutamic Acid and 1% Penicillin-Streptomycin at 37°C in 5% CO2 and transfected with plasmids (Mirus TransIT-X2 Transfection Reagent or Lipofectamine 2000). Imaging was performed 18-24hr following transfection in external Tyrode solution (119mM NaCl, 5mM KCl, 25mM HEPES, 2mM CaCl2, 2mM MgCl2, 33mM Glucose) at pH 7.4. Mouse embryonic fibroblasts (MEFs) were cultured in high-glucose Dulbecco’s Modified Eagle Medium supplemented with 1 mM sodium pyruvate, 1× non-essential amino acids (NEAA; Gibco), 10% fetal bovine serum, and 1% penicillin-streptomycin. Cells were maintained at 37°C in a humidified incubator with 5% CO₂ and passaged every 2–3 days using 0.05% trypsin-EDTA upon reaching ∼80% confluence. For MEFs LysoTracker experiments, 24h after transfection, cells were incubated with LysoTracker Red DND-99 (50 nM) diluted in pre-warmed feeding medium for 20–30 min at 37 °C under light protection. After incubation, cells were washed twice with Tyrode solution to remove excess dye. Live-cell imaging was performed immediately after washing in Tyrode solution. For CLEM experiments, live cell 2pFLIM was performed 2h after Torin application. Cells were plated on a custom-made, gridded coverslip-bottom dishes (based on IWAKI 3922-035; coverslips were affixed with inverse orientation), pre-coated with carbon by vacuum evaporation and subsequently coated with poly-D-lysine (Sigma-Aldrich, P0899), to allow identification of the same cells following fixation.

For in vitro measurements of purified proteins, TOLLES and sREACh-TOLLES proteins were expressed in *Escherichia coli* BL21 cells, with protein induction performed overnight via IPTG at 25°C. Following cell lysis, the proteins were purified via nickel-affinity chromatography using an ÄKTA system. The purified samples were subsequently aliquoted into 96-well plates at a concentration of 0.02 mg/μL. Proteins were incubated in specifically formulated buffer solutions containing a basal salt mixture of 125 mM KCl, 20 mM NaCl, 2 mM CaCl2, and 2 mM MgCl2. The pH 4.0 condition utilized a 25 mM sodium acetate buffer, whereas the pH 7.5 condition utilized a 25 mM HEPES-NaOH buffer ^29^. To assess the effects of acidity and proteolytic activity, the samples were assigned to three experimental groups: (i) pH 7.5 buffer, (ii) pH 4.0 buffer, and (iii) pH 4.0 buffer supplemented with 0.05% (5 units/μL) pepsin. Initial baseline fluorescence lifetime measurements (t=0) were acquired for the pH 4.0 samples using FLIM. Following a 2.5-hour incubation at room temperature, subsequent FLIM measurements were recorded across all three experimental conditions.

Torin (0.5µM), Rapamycin (0.1µM), and Bafilomycin (0.1µM) were purchased from Tocris and incubated overnight, unless noted otherwise, with cells for pharmacological experiments in cell lines. Nigericin/KCl method was used on HEK cells as previously described in ^43^.

### Correlative light and electron microscopy in MEF

Following FLIM imaging, cells were fixed with pre-warmed 2% paraformaldehyde (PFA) and 2.5% glutaraldehyde (GA) in 0.05M phosphate buffer (PB). Cells were incubated with the fixative at 37°C for 1h. Following fixation, cells were washed twice with 1ml of 0.1M PB then incubated overnight in 2ml of 2.5% GA in 0.1M PB. Samples were then washed three times with 1ml of 0.1M PB and maintained in 0.1M PB at 4°C without freezing until further processing for EM up to 4 days later.

Correlative light and electron microscopy was performed essentially as previously described ^76^. Re-imaging of the fixed cells was performed using an A1R confocal microscope (Nikon) equipped with a 100× oil-immersion objective lens (NA 1.45; MRD01905) at 37°C under a 5% CO₂ atmosphere. After washing with 0.1 M phosphate buffer, the cells were post-fixed for 60 min in 0.05 M phosphate buffer containing 1% (w/v) OsO₄ (Electron Microscopy Sciences, 19150) and 1.5% (w/v) potassium ferrocyanide (Kanto Chemical, 32338-30). After rinsing three times with H₂O at RT and three times with H₂O at 50°C, the cells were incubated with Walton’s lead aspartate solution, consisting of 0.635% (w/v) lead nitrate (Sigma-Aldrich, 203580) and 0.4% (w/v) aspartic acid adjusted to pH 5.2 with KOH, for 20 min at 50°C. The cells were dehydrated through an ascending ethanol series: 50% and 70% ethanol for 15 min each on ice, 90% ethanol for 10 min at RT, 95% ethanol for 5 min at RT, and 100% ethanol three times for 5 min each at RT. The dehydrated cells were embedded in Plain Resin (Nisshin-EM) by overlaying the gridded coverslip with resin-filled beam capsules (TAAB). The resin was polymerized at 70°C for 120 h. After polymerization, the gridded coverslip was removed, and the resin block was trimmed to approximately 500 × 500 μm. Serial ultrathin sections, 40 nm in thickness, were cut using an ultramicrotome (EM UC7, Leica) equipped with a diamond knife (Ultra JUMBO, 45°; DiATOME). The sections were collected on cleaned silicon wafer strips and imaged using a scanning electron microscope (JSM-IT800SHL, JEOL). High-magnification imaging was performed in beam deceleration mode using a scintillator backscattered electron detector (SBED), with an accelerating voltage of 1 keV, a specimen bias voltage of 5 keV, a frame size of 5,120 × 3,840 pixels, a working distance of approximately 6.0 mm, a field of view of 25.6 × 19.2 μm, and a dwell time of 14.1 μs. The resulting pixel size was 5 × 5 nm. Image processing, correlation of light and electron microscopy images, three-dimensional visualization, and annotation of autophagic structures were performed using Fiji and Imaris version 9.8.2 (Bitplane). Autophagosomes and autolysosomes were manually identified by visual inspection of the serial EM images based on the ultrastructural criteria described in ^77^. Structures that showed some, but not all, characteristic features of autophagosomes were classified as probable autophagosomes. The numbers of autophagosomes, probable autophagosomes, and autolysosomes were recorded for each region of interest and used to calculate the relative abundance of autolysosomes for comparison with the FLIM measurements.

### C. elegans genetics

*C. elegans* transgenic extrachromosomal array for sR::TOLLES::LGG-1 (*bggEx170*) was generated with the following injection conditions: P*rgef-1*::sR::Tolles::LGG-1 (pBG-GY1117) 2.5ng/µl; P*unc-122*::GFP (pBG-193) 20ng/µl; pBluescript (pBG-49) 77.5ng/µl. Transgenic sR::TOLLES::LGG-1 (*bggEx170*) was mated to *rpm-1 (ju44)* to generate sR::TOLLES::LGG-1; *rpm-1* mutant line.

*C. elegans* were maintained on nematode growth medium agar seeded with OP50 *E. coli*, maintained at 20°C, and 2pFLIM imaged at room temperature. Before experiments, *C. elegans* were synchronized and young adult animals were used for imaging. For imaging, fluorescent positive *C. elegans* were mounted onto a slide with 30 μl M9 buffer and 3 μl Levamisole at 1 μM.

### Immunohistochemistry

Mice were deeply anesthetized by intraperitoneal injection of a high-dose ketamine/xylazine solution, and anesthesia depth was confirmed by the absence of paw pinch withdrawal and corneal reflexes prior to proceeding (an additional half-dose was administered if reflexes persisted). Anesthetized animals were secured to a perfusion platform, and the thoracic cavity was exposed via sequential abdominal, diaphragmatic, and bilateral rib incisions, taking care to avoid damage to the liver, heart, and lungs. A 27G needle connected to a peristaltic pump was inserted into the left ventricle, and the right atrium was incised to allow outflow. Mice were perfused with 15ml PBS followed by 15ml 4%PFA in PBS. Perfused brains were sliced at 80μm using a vibratome (Leica VT1000S) and thereafter stained. Slices were <u>washed</u> three times with PBS for 5 minutes each at room temperature. For Cathepsin D and Atg5 antigens only, antigen retrieval was performed to expose the antigens. Slices were incubated for 20 minutes in pre-warmed citrate buffer (80°C, pH 6.0), followed by cooling at room temperature for 20 minutes. The slices were then washed three times with PBS. For permeabilization, all slices were incubated in PBST containing 1.2% Triton X-100 for 10 minutes. Slices were washed an additional 3 times for 5 minutes each and transferred for 1.5 hours in blocking buffer (5% NGS, 2% BSA, 0.2% Triton in PBS solution) at room temperature. Samples were then incubated in primary antibodies (Anti Parvalbumin, Invitrogen PA1-933, 1:300, Anti p62, 31403-1-AP, 1:200. Anti CathepsinD, Abcam ab75852, 1:100. Anti-Atg5, Abcam 108327, 1:200. anti-WIPI4 – *Wdr45* gene, 19194-1-AP, proteintech, 1:200) diluted in blocking buffer overnight at 4°C. Samples were then washed three times with PBS for 15 minutes and incubated for 60 minutes in secondary Alexa 594 1:1000 (Thermo Fisher Scientific). Slices are washed three times with PBS for 15 minutes each, mounted onto slides, sealed with mounting medium (VectaShield HardSet) and left to dry overnight. Due to inconsistent staining in brain slices for FIP200, we validated efficiency of Cas9/gRNA targeting by co-expressing mouse FIP200-GFP together with the corresponding px330-gRNA or px330-gRNA control in HEK293 cells.

### In Utero Electroporation, AAV injections and cranial windows

In Utero Electroporation was performed on ICR timed pregnant mice at E14.5-15.5 (Envigo). Mice were anesthetized with isoflurane and given 1 μg/g buprenorphine for analgesia. Uterine horns were exposed through an abdominal incision along the *linea alba*, and the lateral ventricle of each embryo was injected with plasmids mixed with 0.01% Fast Green dye (Sigma-Aldrich). Five electrical pulses (40-45V, 50-ms duration, 1 Hz) were delivered using a NEPA21 electroporator (NEPAGENE) with a triple electrode configuration. After birth, fluorescence in the cortex of the pups was visually inspected under a fluorescent lamp. Plasmids were mixed according to experimental conditions described in the results and used at a final concentration of 1µg/µl per plasmid. For sparse labeling, we used AAV injections of a mix of: AAV.pCAG-FLEX-sR-TOLLES-LC3 (1.1×10^13^ vg/ml), AAV9.pCAG-FRT-CyRFP-FRT (5×10^13^ vg/ml) and AAV9.pSyn-FlpO-P2A-Cre (4.2×10^13^ vg/mL, diluted 1:5,000-1:20,000), AAV9.pSyn-XCaMP-R. Injection was performed during the first postnatal week, in p3-p4 mice, which were anesthetized with isoflurane and the right somatosensory cortex was targeted using a glass pipette with a total of 1µl AAV. Pups were then left to recover on a heating pad with their litter and following recovery were returned to their home cage until weaning.

For cranial window surgery, mice (p21-90) were deeply anesthetized with isoflurane for induction (2%–3%) and maintained at surgical plane anesthesia using 1-2% Isoflurane. Mice were administered with 1 μg/g buprenorphine for analgesia and 5 mg/kg carprofen to prevent edema and inflammation. Following fixation in a stereotaxic frame, hair was removed, and the skin and skull were exposed. Then, a 2.3-3mm circular craniotomy was performed over the imaging site using a dental drill. We verified positive fluorescence from IUE/AAV at the somatosensory/motor cortex. The skull was sealed using a 2.3 or 3mm circular cover glass (Matsunami Glass) glued on a 5mm circular cover glass and cemented to skull along with a head-plate to secure the head during imaging using dental cement (C and B Metabond, Parkell). Mice were left to recover and were used for in vivo imaging according to experimental design. Mice that showed signs of window occlusion or tissue damage were excluded from imaging and analysis.

### Correlative light and electron microscopy sample preparation in brain slices

Mice were treated as mentioned earlier (immunohistochemistry) and perfused with 4°C cold flushing solution (0.9% NaCl, 2 mM CaCl₂; 15 mL total, 1.6 mL/min), followed by fixative solution (2% PFA, 2.5% glutaraldehyde in 0.1 M phosphate buffer (PB), pH 7.4, 4°C; 40 mL, 1.6 mL/min). Brains were shipped the following day in PB at 4°C.

Fixed brains were sliced in coronal sections at 200 μm thickness (Leica VT1200 S vibratome). Sections were wet mounted and imaged in 0.1M sodium phosphate buffer in Mattek glass bottom dishes. Images were collected using Zeiss LSM 980 Laser Scanning Confocal with Airyscan. Epifluorescent and brightfield images for location correlation were taken at 10x magnification. Smaller ∼500nm FOVs were imaged at 40x magnification using a silicone immersion objective. Confocal images were taken using ZEISS Airyscan with a 405nm laser at 1.1% laser power. After confocal imaging, samples were processed for TEM. Samples were rinsed three times in 0.1 M sodium cacodylate buffer and post-fixed in 1% osmium tetroxide containing 0.8% K₃[Fe(CN)₆] in 0.1 M sodium cacodylate buffer for 3 h at room temperature, with one solution exchange during incubation. Following postfixation, samples were rinsed with distilled water and stained en bloc with 4% uranyl acetate in ethanol for 2 h. Samples were then dehydrated through a graded ethanol series, infiltrated with Embed-812 resin, and polymerized overnight at 70 °C. Ultrathin sections were cut using a diamond knife (Diatome) on an Enuity ultramicrotome (Leica) and collected onto slot grids. Images were acquired using a JEM-120i transmission electron microscope (JEOL) equipped with a tungsten source operating at 80 kV, and captured using a NanoSprint15 Mk-II (AMT) camera. Correlative light and electron microscopy was then performed to identify the same cells by EM that were imaged by confocal microscopy using inherent fiducial markers in the tissue as previously described ^78^. Lysosomal structures that correlate to the positions of fluorescent puncta were identified by overlaying TEM images with confocal images at matching depths. Correlated puncta/lysosomes were further characterized using ImageJ/Fiji to collect qualitative and quantitative measurements.

### 2pFLIM microscopy

We used a 2pFLIM microscope which was based on a Galvo-Galvo two-photon system (Bergamo, Thorlabs) and a 2pFLIM module (Florida Lifetime Imaging), equipped with a Time-Correlated Single Photon Counting board (Time Harp 260, Picoquant). The microscope was controlled via the FLIMage software. For excitation, we used a Ti:sapphire laser (Chameleon, Coherent) at a wavelength of 860 nm for TOLLES, 920nm for GFP, and 980nm for XCaMP-R/mCherry/mScarlet. Excitation power was adjusted using a Pockels cell (Conoptics) to 1.0–2.0 mW for in vitro experiments and 5-40mW for in vivo experiments. Emission was collected with a 16 × 0.8 NA objective (Nikon), divided with a 565-nm dichroic mirror (Chroma), with emission filters of 525/50 nm and 607/70 nm, and detected with two photomultiplier tubes with low transfer time spread (H7422-40p, Hamamatsu). Images were collected by 128 × 128 or 256 × 256 pixels and acquired at 2 ms/line, averaged over 20-24 frames. For analysis of autophagic vesicle dynamics or XCaMP-R, we imaged continuously at a frame rate of 4Hz for periods of 1-2 minutes and analyzed offline.

### 2pFLIM analysis

All FLIM analysis was performed using a custom C# software (available on the Yasuda lab GitHub: https://GitHub.com/ryoheiyasuda/FLIMage_public). Fluorescence lifetime decay curve A(t) was fitted with a double exponential function convolved with the Gaussian pulse response function:

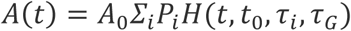

Pi-is the fractional population with the decay time constant of τi, and A0 is the initial fluorescence before convolution. H(t) is an exponential function convolved with the Gaussian instrument response function (IRF).

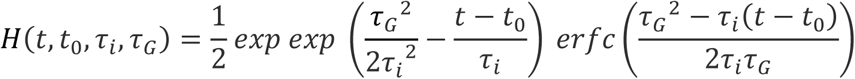

erfc is the error function, τG is the width of the Gaussian pulse response function, and t0 is the time offset. Weighted residuals were calculated using:

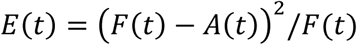

Fitting was performed by minimizing the summed error δ2 = ΣtE(t) for parameters t0, τi (i = 1,2) and τG. We created fluorescence lifetime images by finding the averaged fluorescence lifetime (τm) by the mean photon arrival time subtracted by t0 in each pixel as:

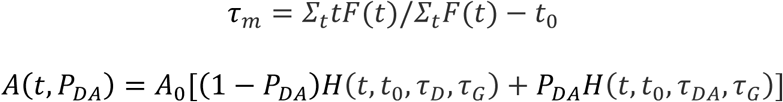

where t0 and τG are obtained from a curve fitting to the fluorescence lifetime decay averaged over all ROIs in an image.

The fluorescence lifetime of TOLLES / sREACh-TOLLES was fixed for TauD/DA values of 3.2/1.4, based on empirical data measured with TOLLES and TOLLES-sREACh pair.

### Puncta detection

Puncta were detected as bright green-channel spots with fluorescence intensity at least 2 standard deviations above the background, located within clearly defined cellular structures identified based on morphology in the red channel, including the soma, dendrites, and axons.

### Motion correction, tracking, and analysis of autophagy vesicles

Single image frames from both channels (green: TOLLES, red: CyRFP) collected simultaneously were used for motion registration. Following previous methods of subcellular registration ^79^ we first corrected image motion of structural images (CyRFP) with rigid and nonrigid methods ^80^ and applied those corrections onto images of TOLLES-labeled puncta. We then selected individual dendritic segments within the same FOV for further analysis, drawing a line segment through selected dendrites and generating a straightened image of both image channels ^81^. We then normalized TOLLES intensity by CyRFP images to isolate autophagy puncta from bright subcellular structures and applied a 0.5-1-micron gaussian filter across single frames. To generate kymograph images, highlighting spatio-temporal movements, we averaged pixel intensity around selected dendrites (width = 5-20 microns) and arranged each frame as a row in the kymograph. Finally, to quantify puncta movements (moving: evident in diagonal lines of kymograph, nonmoving: evident as straight lines), we manually traced individual puncta visible in kymographs for each dendritic segment. Tracked points were used to calculate movement speed in pixels/frame and converted to microns/s to compute total distance traveled for a given movement, total time spent not moving (speed < 0.15 pixels/frame) and quantify changes in direction (speed polarity).

### Statistical analysis

All values are presented as mean ± s.e.m unless otherwise noted. Statistical significance was tested by two-tailed Student’s t test for comparison of two groups, or one way ANOVA followed by post-hoc Tukey’s multiple comparison test for comparison of multiple groups (P < 0.05) using GraphPad Prism 9.0 (GraphPad Software). For all statistical tests * = p < 0.05, ** = p < 0.01, *** = p < 0.001 and **** = p < 0.0001 were considered significant. Sample sizes were not predetermined using statistical methods and were selected based on previous similar experimental design.

## Data Availability

The data supporting the current study are available upon request. Plasmids from this paper will be available through Addgene.

## Code Availability

Software for analysis of the FLIM data is available on the Ryohei Yasuda lab GitHub:

https://GitHub.com/ryoheiyasuda/FLIMage_public.

Software for analysis of autophagy vesicles is available on GitHub:

https://github.com/schollben/autophagy_code.

## Acknowledgments

We would like to thank RIKEN BRC for providing the plasmid encoding TOLLES (RDB18222) through the National Bioresource Project of the MEXT Japan. We thank Zvulun Elazar for generously providing the *Atg5* KO MEF cells. AAVs were produced by the ELSC viral facility in the Hebrew University. We thank Ronen Zaidel Bar, Anat Nizan and Itai Rieger for help with *C. elegans* imaging, Rotem Falach and Roman Klimovich for help with software applications, Avraham Ashkenazi for helpful discussions, Yossi Levi for technical assistance, Yoav Adam for providing the AAV of XCaMP-R, and Klaus Nave for providing the CNP-Cre mice.

We would like to acknowledge the generous support of the Sieratzki Institute (T.L.), the Ben Barres Early Career award from the Chan Zuckerberg Initiative (CZI) (T.L.), the Israel Science Foundation (ISF) grants 1384/21 and 1385/21 (T.L.), and an ERC starting grant number 101040128 (T.L.). We would like to acknowledge the Japan Agency for Medical Research and Development Grant JP19dm0207082 (to YH), Japan Agency for Medical Research and Development Grant JP21wm0525015 (to YH), Japan Science and Technology Agency CREST grant JPMJCR24T6 (YH), and The Asahi Glass Foundation (YH), NIH grant 1S10OD036258-01 (C.T. and A.D.), NIH grant R01NS135763 (B.S.) and Alfred P. Sloan Foundation (B.S.).

## Author contributions

TL conceptualized and supervised the project, performed in vitro and in vivo imaging and analyzed data, MM performed in vitro imaging, animal surgeries, in vivo imaging and analyzed data, KA and YH performed and analyzed CLEM in cell lines, NF, CT, AD, and BS performed and analyzed CLEM in brain slices, YL performed in vivo imaging and analyzed data, GB and BS developed the image analysis pipeline for puncta mobility and analyzed data, NM, SO and TK performed in vitro imaging and assisted with data analysis, MD prepared transgenic worms, MG and SBZ performed purified protein measurements, SE and ML assisted with immunostaining, GU performed in vivo imaging, GN and SS assisted with neuronal cultures, RR and EPB assisted with molecular biology, MM, BG, BS and TL wrote the manuscript with input from all authors.

## Declaration of generative AI and AI-assisted technologies in the manuscript preparation process

During the preparation of this work the authors used ChatGPT and Claude in order to improve grammar. After using this tool/service, the authors reviewed and edited the content as needed and take full responsibility for the content of the published article.

