## Supplemental figures for "In vivo imaging of autophagy dynamics in the intact mammalian brain using a fluorescence lifetime-based sensor"

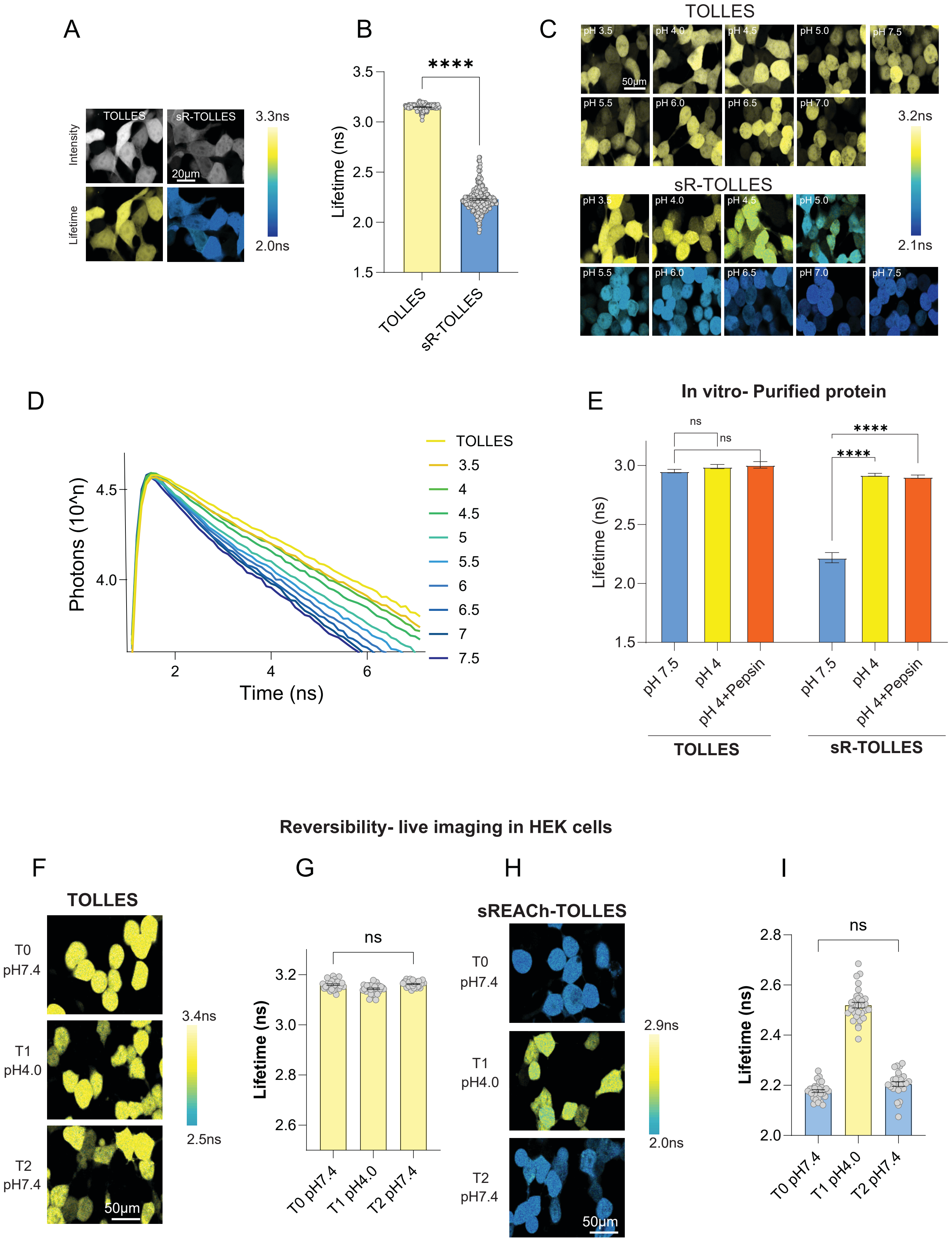
A
TOLLES
sR-TOLLES
Intensity
Lifetime
20µm
3.3ns
2.0ns
B
****
3.5
3.0
2.5
2.0
1.5
Lifetime (ns)
TOLLES
sR-TOLLES
C
TOLLES
pH 3.5
pH 4.0
pH 4.5
pH 5.0
pH 7.5
50µm
pH 5.5
pH 6.0
pH 6.5
pH 7.0
3.2ns
2.1ns
sR-TOLLES
pH 3.5
pH 4.0
pH 4.5
pH 5.0
pH 5.5
pH 6.0
pH 6.5
pH 7.0
pH 7.5
D
Photons (10^n)
4.5
4.0
2
4
6
Time (ns)
TOLLES
3.5
4
4.5
5
5.5
6
6.5
7
7.5
E
In vitro- Purified protein
ns
ns
****
****
Lifetime (ns)
3.0
2.5
2.0
1.5
pH 7.5
pH 4
pH 4+Pepsin
pH 7.5
pH 4
pH 4+Pepsin
TOLLES
sR-TOLLES
Reversibility- live imaging in HEK cells
F
TOLLES
T0
pH7.4
T1
pH4.0
T2
pH7.4
50µm
3.4ns
2.5ns
G
ns
3.2
3.0
2.8
2.6
Lifetime (ns)
T0 pH7.4
T1 pH4.0
T2 pH7.4
H
sREACh-TOLLES
T0
pH7.4
T1
pH4.0
T2
pH7.4
50µm
2.9ns
2.0ns
I
ns
2.8
2.6
2.4
2.2
2.0
Lifetime (ns)
T0 pH7.4
T1 pH4.0
T2 pH7.4

**Extended Fig. 1. In vitro characterization of TOLLES and sR-TOLLES**

**(A)** Representative fluorescence intensity and lifetime images of HEK293 cells expressing TOLLES and sR-TOLLES. Scale bar- 20µm. **(B)** Quantification of mean fluorescence lifetime in cells expressing sR-TOLLES (2.23±0.01ns, n=266 cells) and TOLLES (3.15±0.002ns, n=256 cells). **(C)** Representative lifetime images of TOLLES (top) and sR-TOLLES (bottom) in different pH levels. Scale bar- 50µm. **(D)** Representative fluorescence lifetime curve of HEK293 cells expressing TOLLES and sR-TOLLES in a pH range of 3.5 to 7.5 fitted with a single or double exponential decay, respectively. **(E)** Quantification of mean fluorescence lifetime of purified TOLLES and sR-TOLLES proteins in solutions of pH 7.5, 4.0, and 4.0 supplemented with Pepsin (5 units/µL). TOLLES: pH 7.5: (2.952±0.015, n=6 samples), pH 4: (2.990±0.018, n=5 samples), pH 4+Pepsin: (3.005±0.028, n=5 samples). sR-TOLLES: pH 7.5: (2.218±0.043, n=8 samples), pH 4: (2.921±0.012, n=4 samples), pH 4+Pepsin: (2.905±0.015, n=4 samples). TOLLES: pH 7.5 versus pH 4, p=0.868. pH 7.5 versus pH 4+Pepsin, p=0.67. sR-TOLLES: pH 7.5 versus pH 4 and versus pH 4+Pepsin, $p < 0.0001$. **(F-G)** Representative fluorescence lifetime images **(F)** and quantification **(G)** of HEK293 cells expressing TOLLES in neutral pH (pH 7.4; 3.160±0.003, n=30 cells), following acidification to pH 4 (3.144±0.003, n=34 cells), and after returning to neutral pH (pH 7.4; 3.163±0.002, n=30 cells). p=0.788. **(H-I)** Representative fluorescence lifetime images **(H)** and quantification **(I)** of HEK293 cells expressing sR-TOLLES under neutral pH (pH 7.4; 2.177±0.006, n=29 cells), following acidification to pH 4 (2.520±0.012, n=31 cells), and after returning to neutral pH (pH 7.4; 2.205±0.009, n=28 cells). p=0.098. N=3 biological repetitions. Scale bar- 50µm.

Statistical difference was measured using one-way ANOVA followed by post-hoc Tukey's multiple comparison test (E, G, I) and unpaired two-tailed Student's t-test (B). **** denotes $p < 0.0001$.

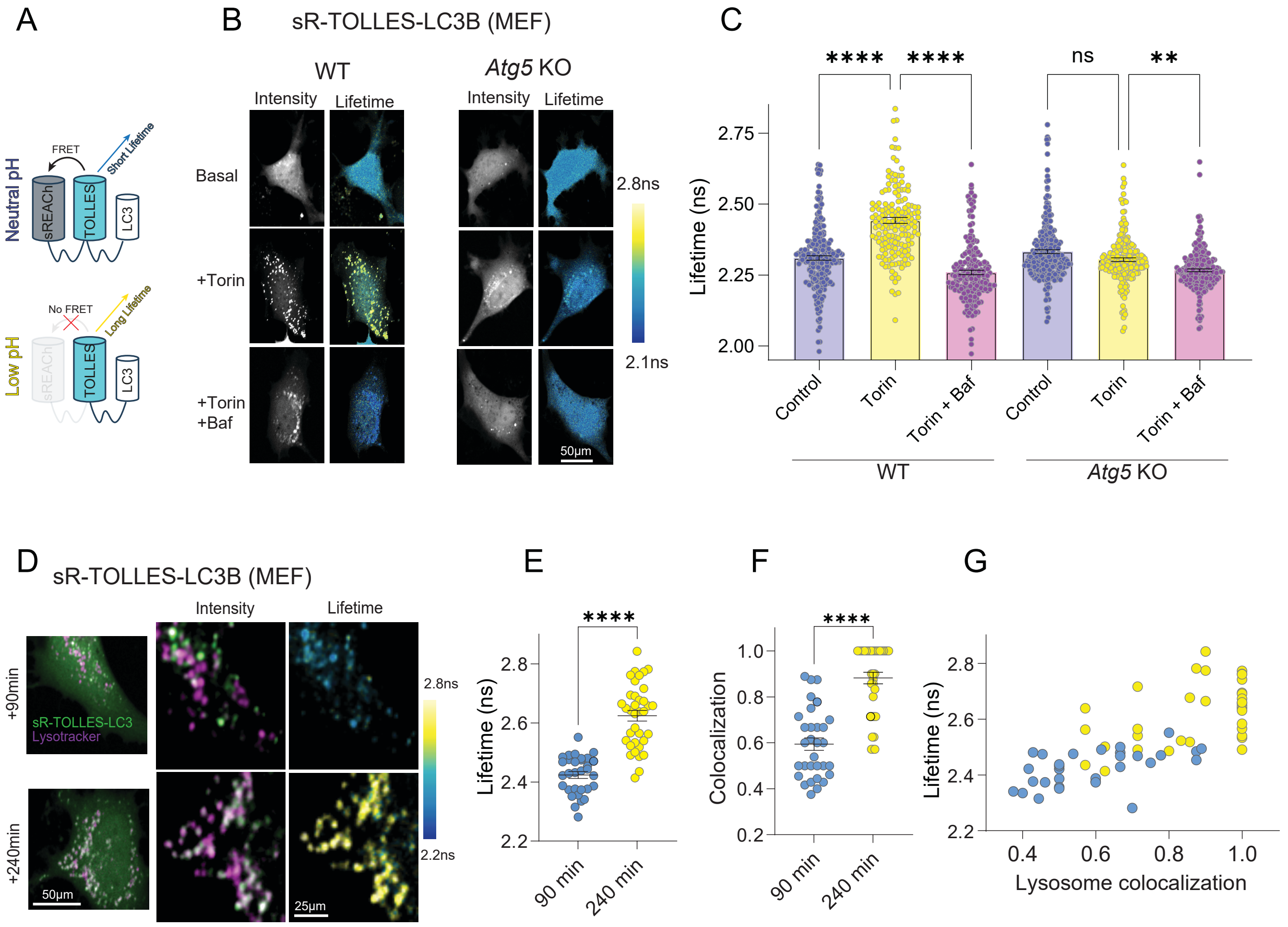
A
Neutral pH
FRET
Short Lifetime
sREACh
TOLLES
LC3
Low pH
No FRET
Long Lifetime
B
sR-TOLLES-LC3B (MEF)
WT
Atg5 KO
Intensity
Lifetime
Basal
+Torin
+Torin
+Baf
2.8ns
2.1ns
50µm
C
****
ns
**
Lifetime (ns)
2.75
2.50
2.25
2.00
Control
Torin
Torin + Baf
WT
Atg5 KO
D
sR-TOLLES-LC3B (MEF)
+90min
+240min
sR-TOLLES-LC3
Lysotracker
50µm
25µm
2.8ns
2.2ns
E
Lifetime (ns)
90 min
240 min
F
Colocalization
G
Lysosome colocalization

**Extended Fig. 2. In vitro validations of lifetime-based autophagy measurements**

**(A)** Schematic illustration of the sR-TOLLES-LC3 sensor in neutral and acidic pH. **(B-C)** Representative intensity and fluorescence lifetime images **(B)** and mean fluorescence lifetime of cells **(C)** in WT and *Atg5* knockout MEFs expressing sR-TOLLES-LC3 under basal conditions, Torin (500nM) and Torin+Bafilomycin (100nM) treatments. (WT: Control: 2.31±0.008ns, n=227 cells; Torin: 2.44±0.010ns, n=168 cells; Baf+Torin: 2.26±0.008ns, n=192 cells. *Atg5* KO: Control: 2.33±0.007ns, n=250 cells; Torin: 2.30±0.006ns, n=201 cells; Baf+Torin: 2.27±0.005ns, n=227 cells). Scale bar- 50µm. **(D)** Representative intensity images of MEF expressing sR-TOLLES-LC3 (green) and Red-Lysotracker (magenta) after 90 and 240 minutes following Torin treatment (left). Higher magnification of the same cells following background subtraction (right). Scale bars are 50µm and 25µm, respectively. **(E)** Quantification of average lifetime of sR-TOLLES-LC3 puncta (90 min: 2.42±0.012ns, n=31 cells, 240 min: 2.62±0.017ns, n=36 cells). **(F)** Fraction of colocalization between TOLLES positive puncta and Lysotracker puncta (90 min: 0.59±0.027, n=31 cells, 240 min: 0.88±0.025, n=36 cells) after Torin treatment. **(G)** Correlation between the lifetime of cellular puncta and colocalization within individual cells. Statistical difference was measured using one-way ANOVA followed by post-hoc Tukey's multiple comparison test (C) and unpaired two-tailed Student's t-test (E-F). **** denotes $p < 0.0001$, ** denotes $p < 0.01$, ns denotes not significant. Atg5 KO Control vs. Torin, p=0.06543. Atg5 KO Torin vs. Torin+Baf, p=0.0039.

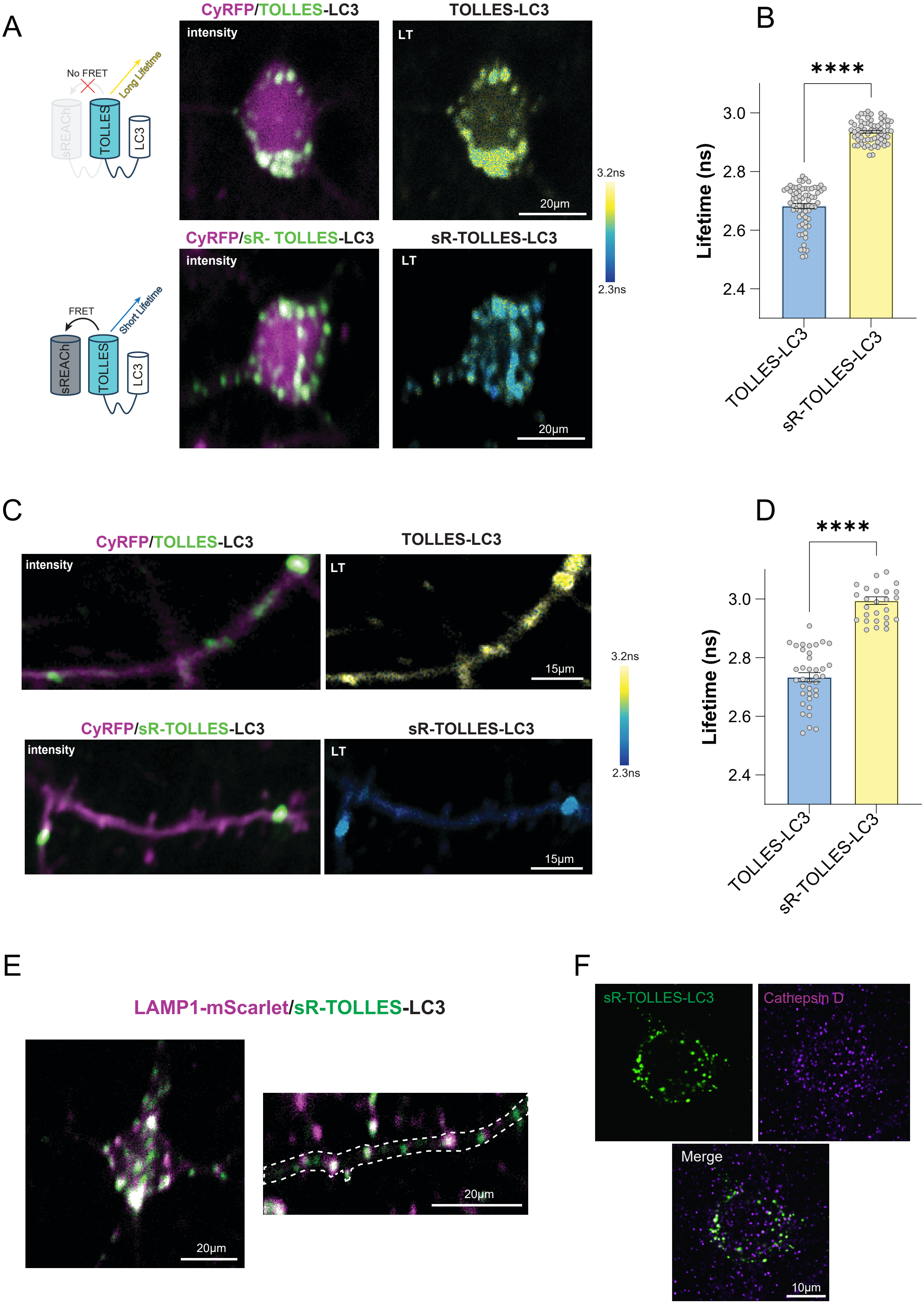

A
No FRET
Long Lifetime
sREACh
TOLLES
LC3
FRET
Short Lifetime
sREACh
TOLLES
LC3
CyRFP/TOLLES-LC3
intensity
TOLLES-LC3
LT
20µm
3.2ns
2.3ns
CyRFP/sR- TOLLES-LC3
intensity
sR-TOLLES-LC3
LT
20µm
B
****
Lifetime (ns)
3.0
2.8
2.6
2.4
TOLLES-LC3
sR-TOLLES-LC3
C
CyRFP/TOLLES-LC3
intensity
TOLLES-LC3
LT
15µm
3.2ns
2.3ns
CyRFP/sR-TOLLES-LC3
intensity
sR-TOLLES-LC3
LT
15µm
D
****
Lifetime (ns)
3.0
2.8
2.6
2.4
TOLLES-LC3
sR-TOLLES-LC3
E
LAMP1-mScarlet/sR-TOLLES-LC3
20µm
20µm
F
sR-TOLLES-LC3
Cathepsin D
Merge
10µm

**Extended Fig. 3. Control experiments for in vivo 2pFLIM of autophagy**

**(A)** *In vivo* 2pFLIM representative images of fluorescence intensity (left) and lifetime (right) of L2/3 neurons expressing CyRFP (magenta), and TOLLES-LC3 (top, green), or sR-TOLLES-LC3 (bottom). Scale bar- 20µm. **(B)** Quantification of mean fluorescence lifetime of L2/3 soma expressing TOLLES-LC3 (2.93±0.008ns, n=68 cells, 3 mice) and sR-TOLLES-LC3 (2.68±0.005ns, n=65 cells, 3 mice). **(C)** Representative fluorescence intensity and lifetime images of L2/3 dendrites expressing TOLLES-LC3 (top) and sR-TOLLES-LC3 (bottom). Scale bars- 15µm. **(D)** Quantification of mean fluorescence lifetime of LC3 positive puncta in dendrites expressing TOLLES-LC3 (2.99±0.013ns, n=29 puncta, 3 mice) and sR-TOLLES-LC3 (2.73±0.015ns, n=37 puncta, 3 mice). **(E)** Representative fluorescence intensity image of a L2/3 neuron co-expressing LAMP1-mScarlet (magenta) and sR-TOLLES-LC3 (green). Pearson correlation coefficient for colocalization analysis for in vivo expression of LAMP1 and sR-TOLLES-LC3 (R=0.872, n=18 cells, 3 mice). Scale bars- 20µm. **(F)** Representative image of soma in fixed brain tissue expressing sR-TOLLES-LC3 (green) and stained with Cathepsin-D antibody (magenta). Colocalization rate of Cathepsin D on TOLLES puncta (45% of sR-TOLLES puncta colocalized with CTSD puncta, n=37 cells, N=2 mice). Scale bar- 10µm. Statistical difference was measured using unpaired two-tailed Student's t-test. **** denotes $p < 0.0001$.

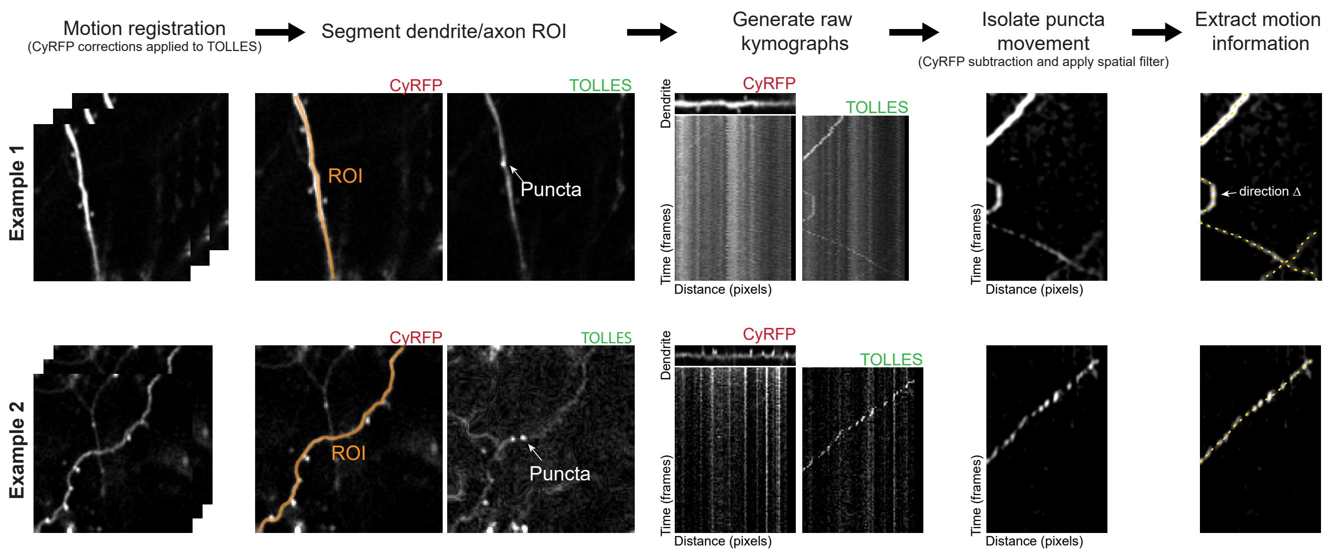

Motion registration
(CyRFP corrections applied to TOLLES)
Segment dendrite/axon ROI
Generate raw kymographs
Isolate puncta movement
(CyRFP subtraction and apply spatial filter)
Extract motion information
Example 1
Example 2
CyRFP
TOLLES
ROI
Puncta
Dendrite
Time (frames)
Distance (pixels)
direction Δ

**Extended Fig. 4. Image processing and analysis pipeline for in vivo autophagic vesicle movement**

Two example fields of view (FOVs; top and bottom) from two-photon images collected from the red (CyRFP/structure) and green (TOLLES, autophagy sensor) channels are first motion corrected. Motion correction is performed on structural images and applied to autophagy sensor images. Individual dendrites within the FOV are then segmented to generated straightened images and subsequent kymographic images. Raw kymographs of the TOLLES sensor are processed further: bleed-through from CyRFP is removed and a spatial Gaussian filter (1 micron sigma) is applied. The resulting kymographs are then used to track individual moving puncta. Track points span each kymographic trajectory and are extracted to calculate motion statistics (velocity, movement distance, changes in direction, etc.).

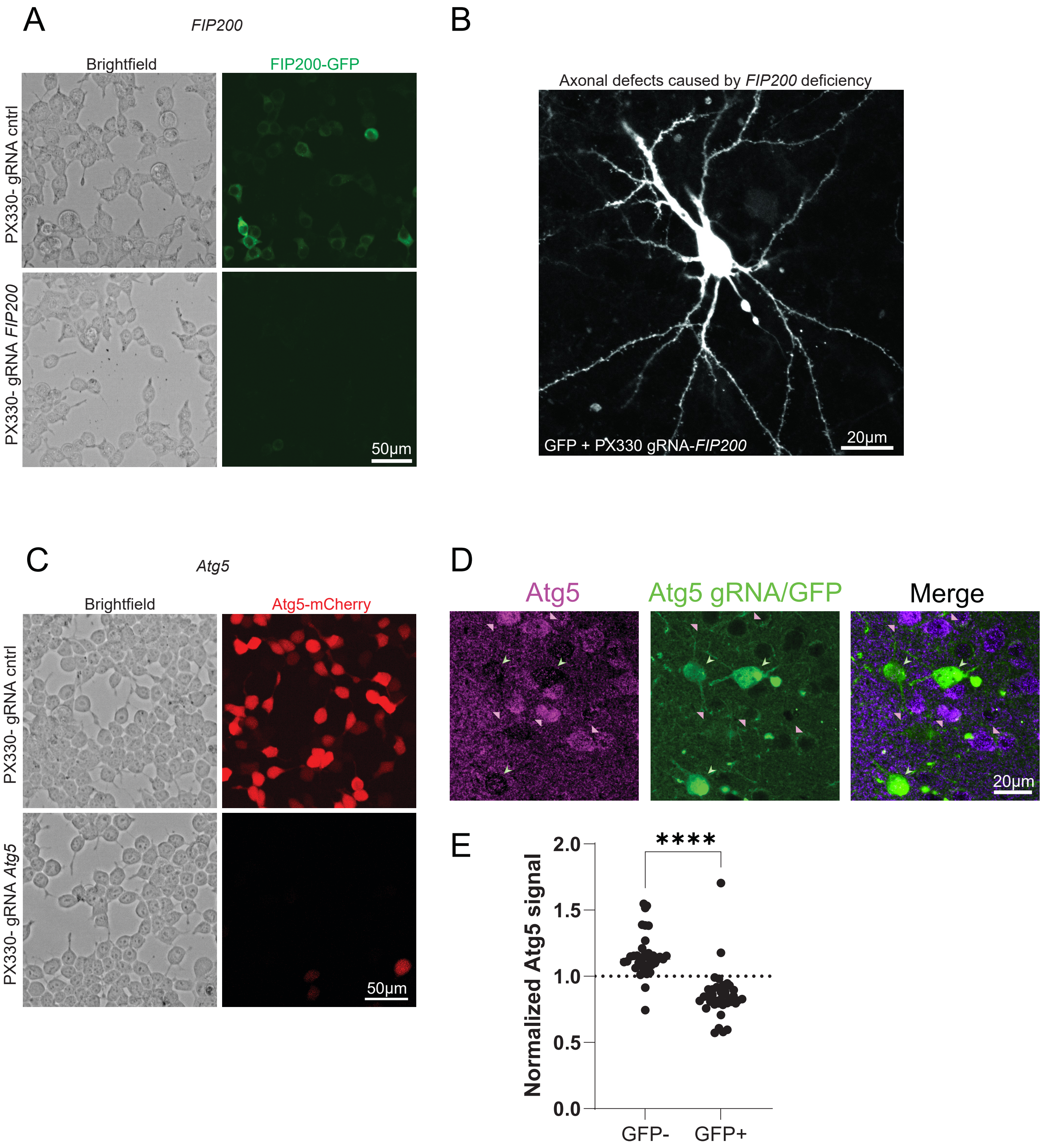

A
FIP200
Brightfield
FIP200-GFP
PX330- gRNA cntrl
PX330- gRNA FIP200
50μm
B
Axonal defects caused by FIP200 deficiency
GFP + PX330 gRNA-FIP200
20μm
C
Atg5
Brightfield
Atg5-mCherry
PX330- gRNA cntrl
PX330- gRNA Atg5
50μm
D
Atg5
Atg5 gRNA/GFP
Merge
20μm
E
Normalized Atg5 signal
2.0
1.5
1.0
0.5
0.0
****
GFP-
GFP+

**Extended Fig. 5. Validation of CRISPR/Cas9-based targeting of *Atg5* and *FIP200***

**(A)** Representative bright-field and fluorescence images of HEK cells transfected with FIP200-GFP and either Cas9/control gRNA (top) or *FIP200* gRNA (bottom), using the same plasmids as those used *in vivo* for *FIP200*. Scale bar- 50µm. **(B)** Representative image of cortical neurons expressing GFP/Cas9/gRNA *FIP200*, showing axonal swelling phenotype. Scale bar- 20µm. **(C)** Representative bright-field and fluorescence images of HEK cells transfected with Atg5-mCherry and either Cas9/control gRNA (top) or *Atg5* gRNA (bottom), using the same plasmids as those used *in vivo* for *Atg5* KO. Scale bar- 50µm. **(D)** Representative image of fixed brain slice, with cortical neurons expressing GFP (green) and Cas9/gRNA targeting *Atg5*, co-stained for Atg5 (magenta). Arrowheads indicate GFP-positive (green) and nearby non-expressing (pink) cells. Scale bar- 20µm. **(E)** Quantification of normalized Atg5 signal in GFP/Cas9/gRNA Atg5-positive cells (GFP+; 0.855±0.030, n=37 cells) compared with nearby non-expressing cells (GFP-; 1.157±0.027, n=37 cells).
Statistical difference was measured using an unpaired two-tailed Student's *t*-test. **** denotes p < 0.0001.

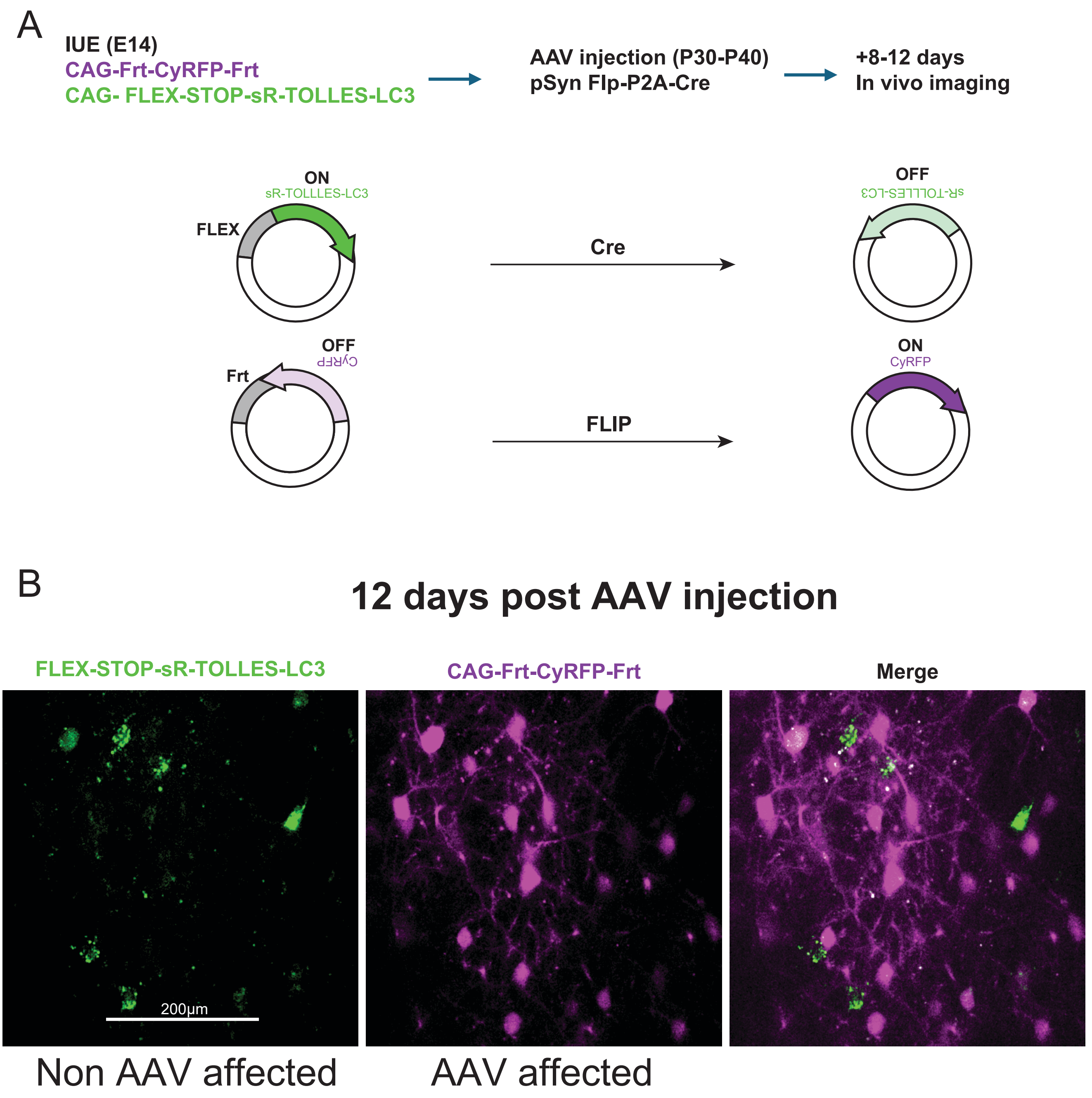

A
IUE (E14)
CAG-Frt-CyRFP-Frt
CAG- FLEX-STOP-sR-TOLLES-LC3
AAV injection (P30-P40)
pSyn Flp-P2A-Cre
+8-12 days
In vivo imaging
ON
sR-TOLLLES-LC3
FLEX
Cre
OFF
sR-TOLLES-LC3
OFF
CyRFP
Frt
FLIP
ON
CyRFP
B
12 days post AAV injection
FLEX-STOP-sR-TOLLES-LC3
CAG-Frt-CyRFP-Frt
Merge
200μm
Non AAV affected
AAV affected

**Extended Fig. 6. In vivo turnover of sR-TOLLES-LC3 following Cre-mediated expression shutoff**

**(A)** Scheme of the FLApHY Stop experiment. In utero electroporation (E14) was used to deliver plasmids encoding FLEX-STOP-sR-TOLLES-LC3 and Frt-CyRFP (left). At P30–40, mice were injected with AAV encoding pSyn-Flp-P2A-Cre (middle). 8-12 days following AAV injection, in vivo imaging was performed to assess expression (right). **(B)** Representative images acquired 12 days after AAV injection, showing fields of view of cells that were not infected by the virus and express sR-TOLLES-LC3 (left, green), and cells that were infected and no longer expressing sR-TOLLES-LC3, now expressing CyRFP (middle, magenta), together with the merged image (right). Scale bar- 200μm.

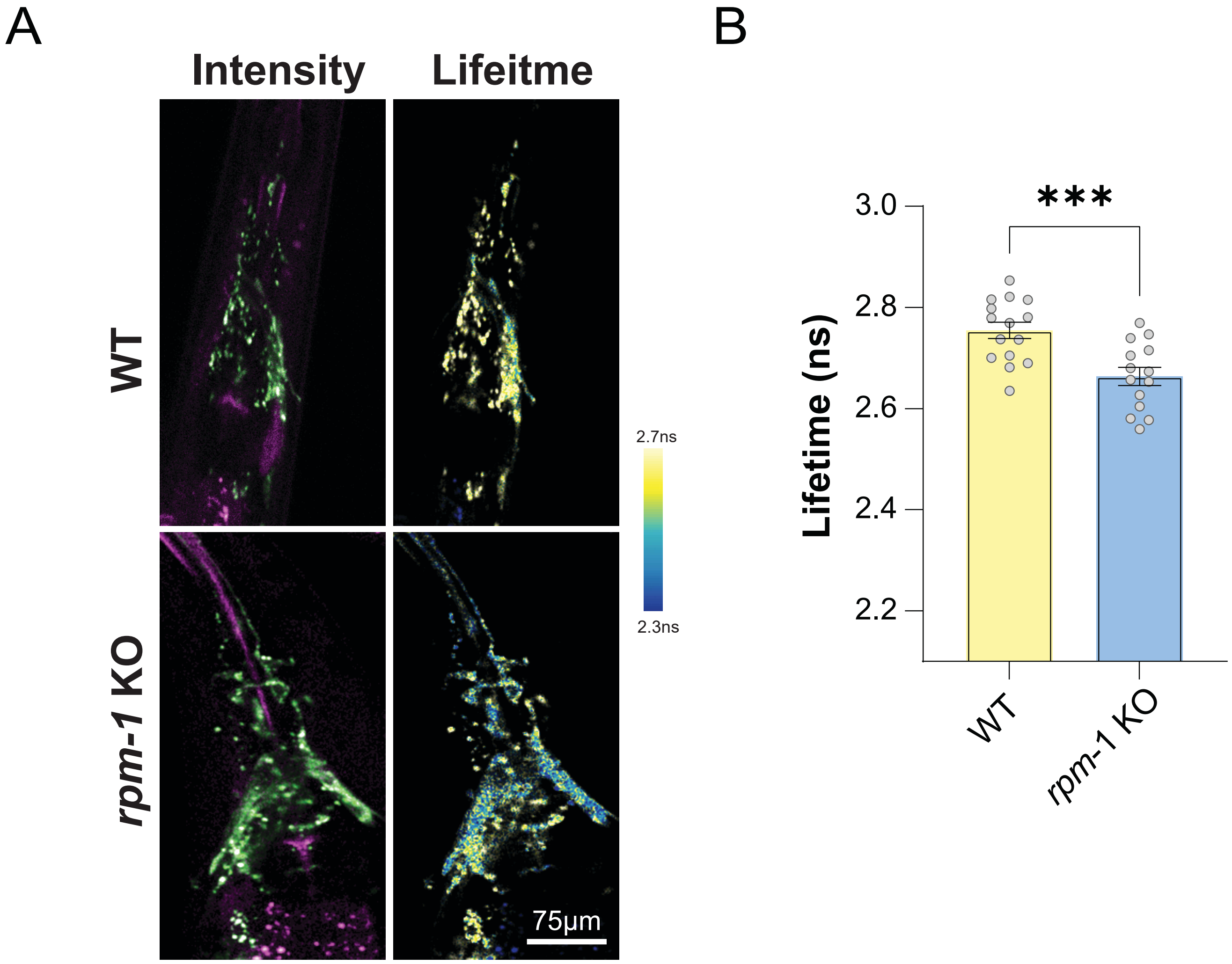
A
Intensity
Lifeitme
WT
rpm-1 KO
2.7ns
2.3ns
75μm
B
***
3.0
2.8
2.6
2.4
2.2
Lifetime (ns)
WT
rpm-1 KO

**Extended Fig. 7. Autophagy measurements in *C. elegans* expressing sR::TOLLES::LGG-1**
**(A)** Representative fluorescence intensity (left) and lifetime (right) images of transgenic *C. elegans* expressing sR::TOLLES::LGG-1 in WT (top) or *rpm-1* mutant. Scale bar- 75µm. **(B)** Quantification of fluorescence lifetime for WT (2.75±0.016ns, n=15) and *rpm-1* mutants (2.66±0.018ns, n=14) in the axonal bundle that forms the nerve ring, p=0.0008. Note that reduced fluorescence lifetime indicates increased formation of autophagosomes. Statistical difference was measured using unpaired two-tailed Student's t-test. *** denotes $p < 0.001$.

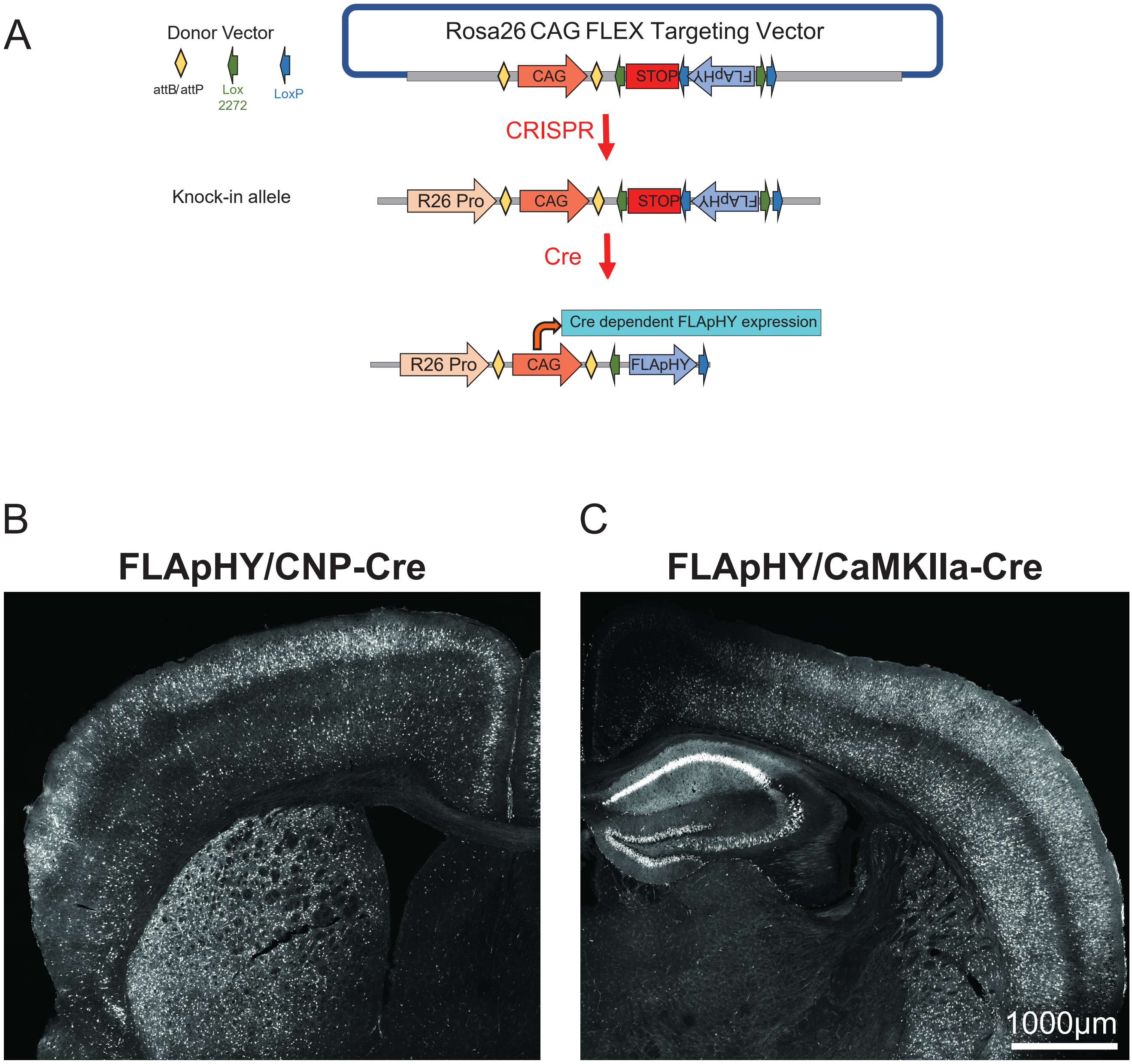
A
Donor Vector
attB/attP
Lox 2272
LoxP
Rosa26 CAG FLEX Targeting Vector
CAG
STOP
FLApHY
CRISPR
Knock-in allele
R26 Pro
CAG
STOP
FLApHY
Cre
Cre dependent FLApHY expression
R26 Pro
CAG
FLApHY
B
FLApHY/CNP-Cre
C
FLApHY/CaMKIIa-Cre
1000µm

**Extended Fig. 8. sR-TOLLES-LC3 Knock-In transgenic mouse description**

(**A**) Schematic illustrating the generation of the FLApHY knock-in (KI) mouse line and the genomic insertion strategy for Cre-dependent expression of the FLApHY transgene from the ROSA26 (R26) locus.

(**B-C**) Representative images showing FLApHY expression across the cerebral hemisphere in FLApHY KI mice crossed with either CNP-Cre (**B**) or CaMKIIα-Cre (**C**) mice, enabling cell-type-specific expression in oligodendrocytes or excitatory neurons, respectively. Scale bar- 1,000µm.

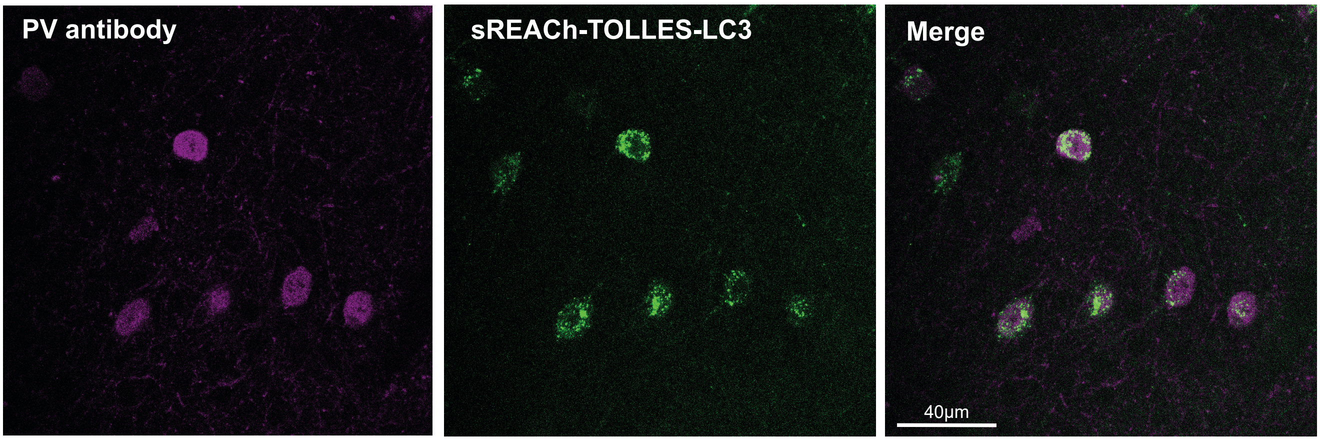
PV antibody
sREACh-TOLLES-LC3
Merge
40μm

**Extended Fig. 9. Validation of PV cells targeting of sR-TOLLES-LC3**

Representative images of fixed brain slices from a PV-Cre transgenic mouse expressing AAV-pCAG-FLEX-sR-TOLLES-LC3. PV antibody (magenta), sR-TOLLES-LC3 (green), and the merged image. Scale bar- 40µm.

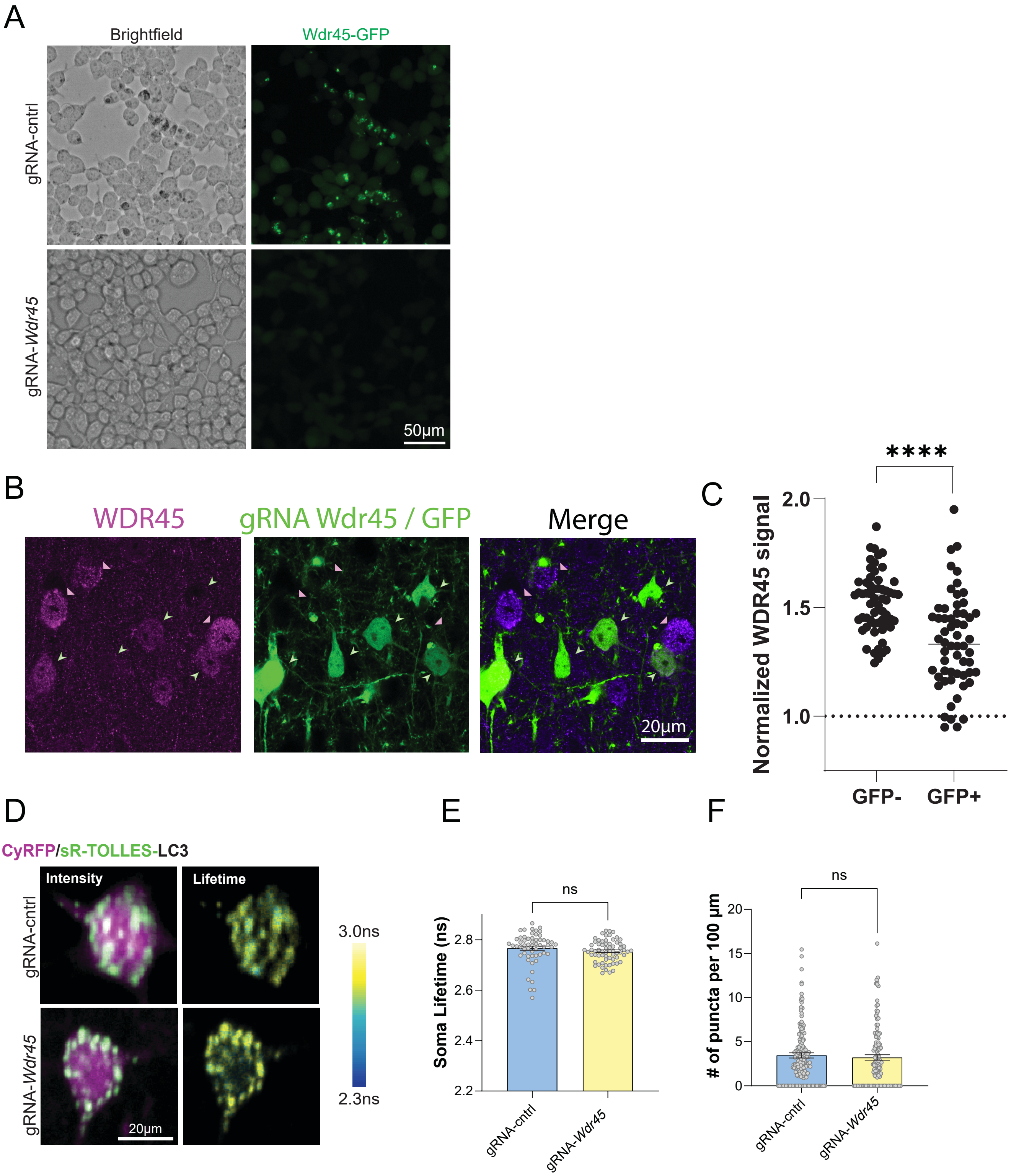

A
Brightfield
Wdr45-GFP
gRNA-cntrl
gRNA-Wdr45
50µm
B
WDR45
gRNA Wdr45 / GFP
Merge
20µm
C
****
Normalized WDR45 signal
2.0
1.5
1.0
GFP-
GFP+
D
CyRFP/sR-TOLLES-LC3
Intensity
Lifetime
gRNA-cntrl
gRNA-Wdr45
20µm
3.0ns
2.3ns
E
ns
Soma Lifetime (ns)
2.8
2.6
2.4
2.2
gRNA-cntrl
gRNA-Wdr45
F
ns
### of puncta per 100 µm
20
15
10
5
0
gRNA-cntrl
gRNA-Wdr45

**Extended Fig. 10. *WDR45* validation and additional data**

(**A**) Representative bright-field and fluorescence images of HEK cells transfected with WDR45-GFP and either CRISPR/Cas9 gRNA-control (top) or gRNA-*WDR45* (bottom), using the same plasmid as that used *in vivo*. Scale bar- 50µm. (**B**) Representative image of cortical neurons expressing GFP (green) and CRISPR/Cas9-gRNA *WDR45*, stained for WDR45 (magenta). Arrowheads indicate GFP-positive (green) and nearby non-expressing (pink) cells. Scale bar- 20µm. (**C**) Quantification of normalized WDR45 staining in cells expressing GFP/Cas9/gRNA-*WDR45*, compared with nearby non-expressing cells (GFP+: 1.343±0.028, n=58 cells; GFP-: 1.563±0.027, n=63 cells). (**D**) Representative fluorescence intensity and lifetime images of cells expressing sR-TOLLES-LC3 (green) and CyRFP (magenta) alongside Cas9/gRNA-control (top) and gRNA-*WDR45* (bottom). Scale bar- 20µm. (**E**) Mean fluorescence lifetime in soma of Cas9/gRNA-*WDR45* (2.755±0.005ns, n=65 cells) and gRNA-control (2.767±0.008ns, n=57 cells) cells. p=0.2152. N=4-5 mice. (**F**) Quantification of the number of LC3-positive puncta per 100µm of dendrite in Cas9/gRNA-*WDR45* expressing (3.225±0.308 puncta/100µm, n=125 dendritic branches) and gRNA-control (3.457±0.293 puncta/100µm, n=165 dendritic branches) neurons (p=0.589). N=4 mice.

Statistical difference was measured using unpaired two-tailed Student's t-test (C, E, F). **** denotes p < 0.0001.

| Cell | Puncta | Size (Diameter in µm) | Mean intensity | Content (no. of electron dense items) | Image |
|---|---|---|---|---|---|
| 1 | 1 | 0.298 | 128.665 | 2 | 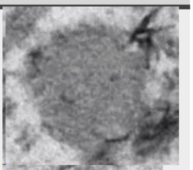 |
| 1 | 3 | 0.349 | 107.342 | 0 | 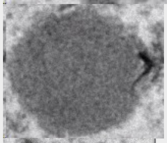 |
| 1 | 4 | 0.437 | 111.803 | 0 | 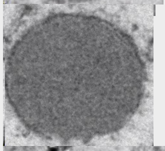 |
| 1 | 5 | 0.425 | 132.437 | 1 | 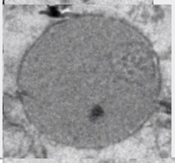 |
| 3 | 1 | 0.197 | 131.701 | 0 | 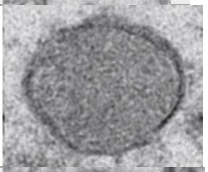 |
| 4 | 1 | 0.394 | 106.273 | 1 | 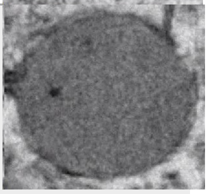 |
| 4 | 2 | 0.337 | 108.697 | 1 | 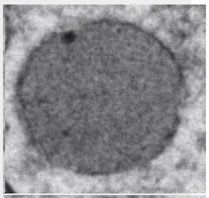 |
| 4 | 3 | 0.275 | 117.146 | 1 | 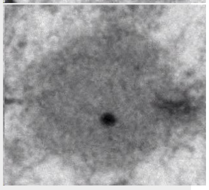 |
| 4 | 4 | 0.339 | 117.82 | 3 | 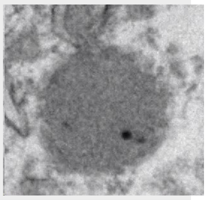 |
| 5 | 1 | 0.348 | 115.966 | 2 | 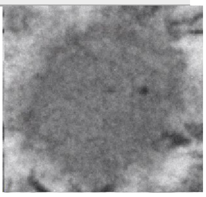 |

| | | | | | |
|---|---|---|---|---|---|
| 5 | 2 | 0.471 | 97.104 | 0 | 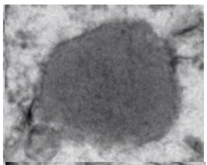 |
| 5 | 4 | 0.543 | 114.811 | 2 | 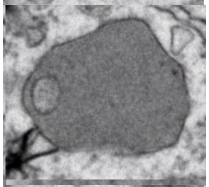 |
| 5 | 7 | 0.403 | 133.853 | 0 | 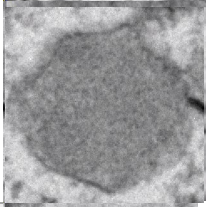 |
| 6 | 1 | 0.261 | 114.843 | 0 | 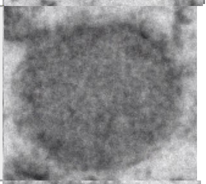 |
| 7 | 1 | 0.367 | 123.97 | 0 | 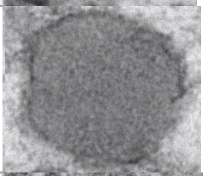 |
| 8 | 1 | 0.41 | 110.452 | 0 | 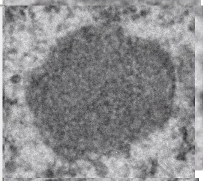 |
| 8 | 2 | 0.598 | 123.015 | 2 | 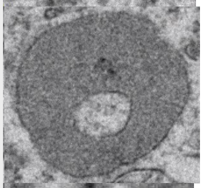 |
| 8 | 3 | 0.209 | 109.968 | 0 | 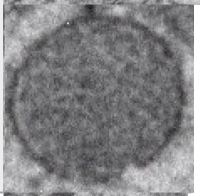 |
| 9 | 1 | 0.458 | 97.361 | 1 | 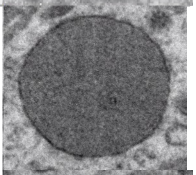 |
| 9 | 2 | 0.32 | 94.659 | 1 | 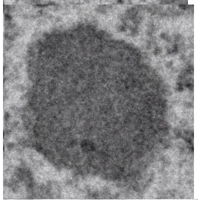 |

| | | | | | |
|---|---|---|---|---|---|
| 9 | 3 | 0.54 | 100.953 | 3 | 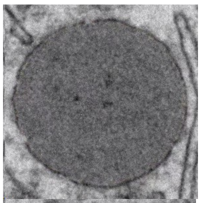 |
| 9 | 4 | 0.4 | 106.284 | 0 | 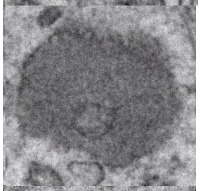 |
| 9 | 5 | 0.349 | 119.116 | 1 | 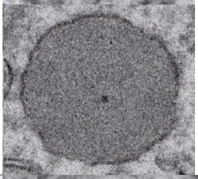 |
| 10 | 2 | 0.444 | 95.524 | 1 | 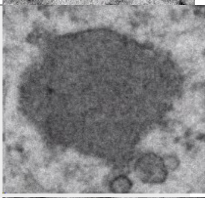 |
| 10 | 3 | 0.169 | 106.55 | 1 | 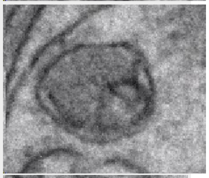 |
| 10 | 4 | 0.299 | 102.597 | 1 | 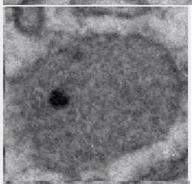 |
| 10 | 5 | 0.445 | 68.075 | 0 | 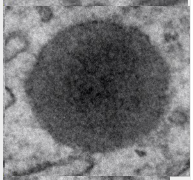 |
| 10 | 6 | 0.339 | 95.275 | 2 | 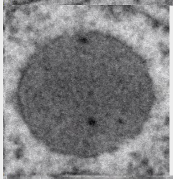 |
| 10 | 7 | 0.263 | 107.47 | 1 | 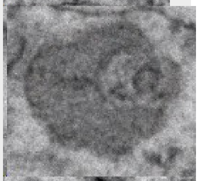 |
| 10 | 8 | 0.439 | 105.621 | 1 | 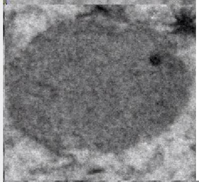 |
| 10 | 9 | 0.577 | 109.374 | 0 | 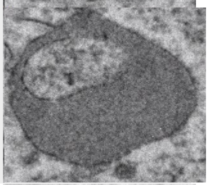 |

| | | | | | |
|---|---|---|---|---|---|
| 10 | 11 | 0.376 | 108.18 | 0 | 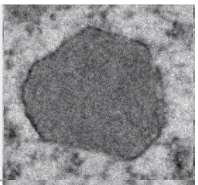 |
| 11 | 1 | 0.322 | 105.439 | 1 | 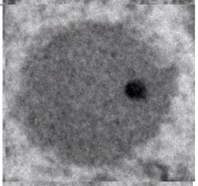 |
| 11 | 3 | 0.44 | 109.091 | 4 | 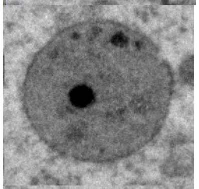 |
| 11 | 4 | 0.331 | 111.912 | 0 | 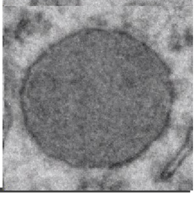 |
